# Rodent Origins and Human-Mediated Evolutionary Dynamics of PRRSV

**DOI:** 10.1101/2025.11.25.690366

**Authors:** Dinko Novosel, Andreja Jungic, Omar Rota-Stabelli, Marti Cortey, Ino Curik, Enric Mateu

## Abstract

Porcine reproductive and respiratory syndrome virus (PRRSV) is among the most economically damaging swine pathogens worldwide, yet its evolutionary origins and the forces driving its diversification remain unclear. Using complete arterivirus genomes, we applied an integrative, high-resolution phylogenomic framework that combined time-calibrated evolutionary inference with recombination- and introgression-aware analyses. This approach resolved the divergence history of PRRSV and uncovered a cryptic mosaic genome and reticulation shaped by recurrent recombination and interspecies viral gene flow. We show that North American and European lineages diverged in the late 19th century – long before the emergence of disease on either continent and that their closest relatives are rodent arteriviruses, indicating a rodent ancestry. Genome-wide scans further reveal that PRRSV evolution is driven by adaptive changes in the glycoprotein ORF3, whereas the replicase genes remain a conserved genomic backbone. Demographic reconstructions suggest that the intensification of pig farming in the mid-20th century-imposed transmission bottlenecks that amplified genetic drift and accelerated the fixation of advantageous variants. Together, these findings rewrite the evolutionary timeline of PRRSV and provide a framework for understanding how human practices shape the emergence and spread of viral pathogens across animal and human health.

## Introduction

First identified in the late 1980s in North America ^1,2^, porcine reproductive and respiratory syndrome (PRRS) has since spread worldwide and remains one of the most economically devastating diseases in the swine industry ^3,4^. Almost simultaneously, a similar syndrome emerged in Europe ^5^. Researchers soon determined that the European and North American outbreaks were caused by two related but genetically distinct viruses ^6,7^. These strains were initially classified as Type 1 (European) and Type 2 (North American) PRRS viruses and are now recognised as two separate species (PRRSV-1 and PRRSV-2) within the genus *Betaarterivirus* in the family *Arteriviridae*. PRRSV genome is a positive-sense single-stranded RNA encoding at least ten open reading frames (ORFs) ^8–10^. Two of these ORFs (1a and 1b) encode the non structural proteins including the replication machinery. Whereas the remaining ORFs (2-7) encode the structural proteins ^11^.PRRSV is genetically and antigenically diverse ^12,13^, which has hindered efforts to trace its origin and early evolution. Early phylogenetic analyses suggested that the European PRRSV lineage emerged from a North American source ^14,15^. However, a later study found that strains circulating in Eastern Europe exhibited greater genetic diversity than those in North America, indicating to Eastern Europe as a possible origin ^16^. Estimates of the timing of PRRSV divergence have also varied widely. One molecular clock analysis placed the virus’s most recent common ancestor around 1979 ^17^. In contrast, a study using larger datasets traced the divergence of the two PRRSV species back to the late 19^th^ century ^18^. Another investigation estimated that PRRSV-1 began diversifying between 1946 and 1967, whereas PRRSV-2 began diversifying between 1977 and 1981 ^19^. Together, these differing findings highlight the uncertainty surrounding when and where PRRSV first emerged. Reconstructing the evolutionary history of a fast-evolving RNA virus like PRRSV (whose substitution rate has been estimated at approximately ∼1×10^-3^ substitutions sites^-1^ year^-1^) is analytically challenging. Conventional phylogenetic methods such as neighbour-joining or standard maximum likelihood often perform poorly for such rapidly evolving pathogens because they do not account for the time-dependent nature of sequence change. Bayesian phylogenetic frameworks, which incorporate molecular clock models and coalescent theory, have greatly improved time-calibrated virus phylogenies.

PRRSV exemplifies the evolutionary dynamics of RNA viruses: most mutations are eliminated by purifying selection, while rare advantageous changes particularly in the immunodominant glycoprotein GP5—are fixed by diversifying selection to promote immune escape ^20^. Adaptive changes cluster in surface glycoproteins, whereas replicative and internal proteins remain conserved. Frequent homologous recombination among co-circulating strains produces mosaic genomes fuelling the emergence of novel variants with enhanced fitness.

Despite the attention the PRRS virus has received, due to its economic importance and widespread distribution, many aspects of its origin and evolution remain unresolved. These include the timing of divergence between PRRSV-1 and PRRSV-2, the distribution of selection pressures across the genome, and the potential role of recombination– including introgression from rodent arteriviruses– in shaping viral origins ^21^. The recent discovery of diverse rodent arteriviruses closely related to PRRSV has intensified questions about its ancestral reservoir and raised the possibility that cross-species transmission shaped its origins^22^.

Moreover, PRRSV serves as a model for how anthropogenic changes in livestock production can accelerate viral emergence and evolution. Here, we use a holistic phylogenomic approach to complete arterivirus genomes, combining Bayesian time-calibrated inference, ancestral recombination graph reconstruction, species-tree reconciliation, and genome-wide introgression and selection analyses. This strategy allows us to distinguish divergence from recombination, identify adaptive changes in viral proteins, and link demographic dynamics to shifts in pig husbandry. Together, these analyses establish a framework for resolving the origins and diversification of PRRSV and provide a general model for studying the emergence of animal and human pathogens.

## Results

### Rodent *arteriviruses* are the closest known relatives of PRRSV

Whole-genome phylogeny (Fig. 1a) reveals pronounced genetic structure across *Arteriviridae*. Within *Betaarterivirus*, PRRSV-1 and PRRSV-2 form reciprocally monophyletic, long-branched clades separated by deep internal nodes. Rodent arteriviruses – rat arterivirus, LDEV, and African pouched rat arterivirus – are the closest sampled relatives of PRRSV, while equine and simian arteriviruses each form distinct clades. Tip density indicates sampling asymmetry, with many more PRRSV genomes from North America than from Europe, reflecting greater sequencing effort in the Americas. Time-calibrated whole-genome phylogenies provide high-resolution insight into the divergence of PRRSV-1 and PRRSV-2, which emerge as two deeply divergent species within *Betaarterivirus* (Fig. 1a-b, Suppl. Fig. 1–2). The historical annotations highlight the intersection of viral evolution with changes in pig farming that likely influenced transmission dynamics. Molecular dating consistently places their most recent common ancestor in the late 19th century (95% HPD: 1870–1905), decades before the first recognised outbreaks in the 1980s and prior to the intensification of industrial pig production. Analyses based on complete genomes identify rodent arteriviruses (RAV, LDEV, APRAV) as the closest relatives to PRRSV, with equine arterivirus (EAV) more distant (Fig. 1b, Suppl. Fig. 1–2).

**Figure 1.**
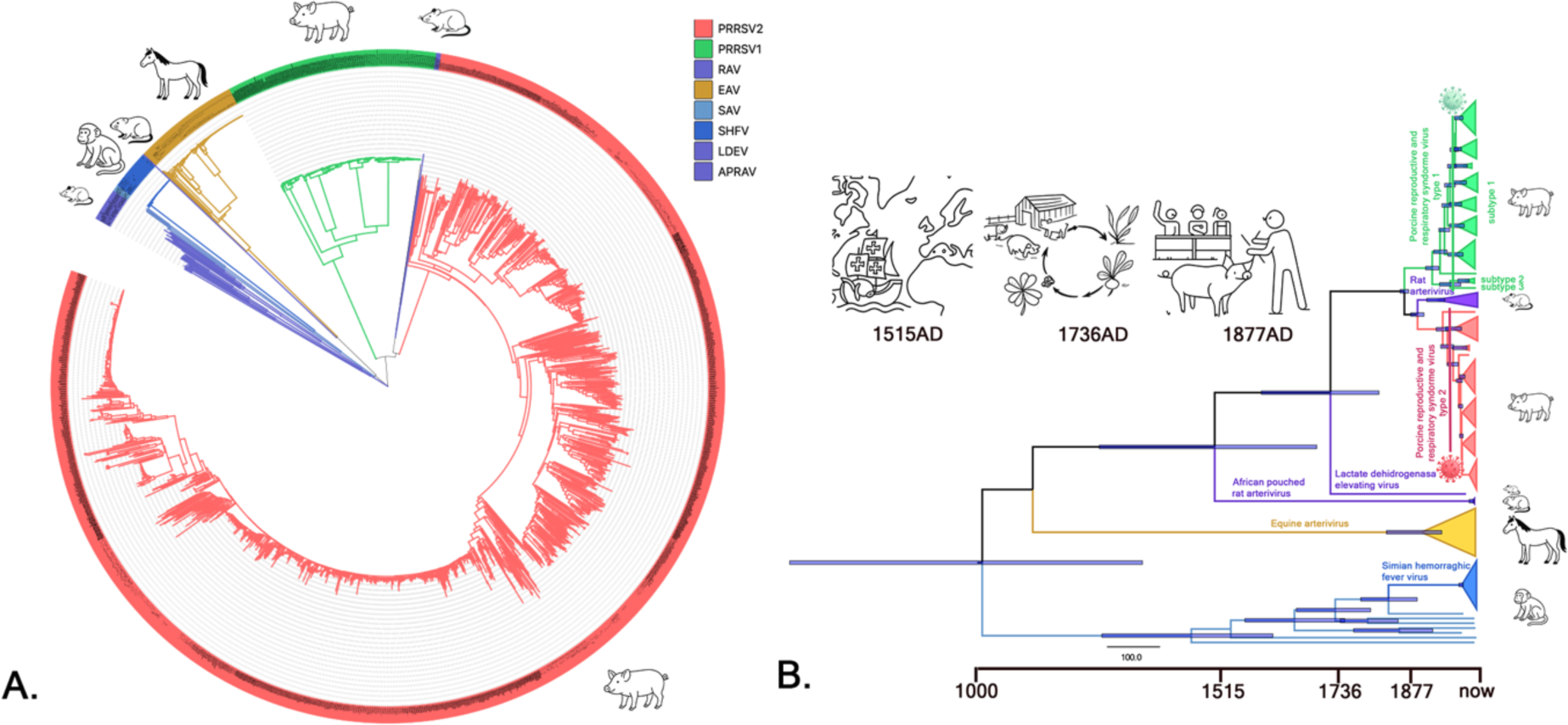
Global diversity and deep-time evolution of PRRSV within *Arteriviridae*. A) Circular whole-genome phylogeny of 1,818 arterivirus genomes. B) Time-calibrated Bayesian phylogeny inferred from a subsample of 115 whole-genome arterivirus. Node bars are the 95% HPD intervals. Clades corresponding to PRRSV-2, PRRSV-1, RAV, EAV, SHFV, LDEV, and APRAV are coloured according to legend. Drawings indicate historical events in swine production and PRRS outbreaks.

Across relaxed-clock models, the best-supported model estimate (Suppl. fig. 2) for the whole genome was 2.38 × 10⁻³ substitutions·site⁻¹·year⁻¹, with divergence times robust to model choice and data curation. Introgression tests revealed a reticulate component in PRRSV evolution. Patterson’s D and the f₄-ratio detected significant gene flow between PRRSV and rodent arteriviruses, with the strongest signals in ORF1a, ORF4, and ORF5 (Fig. 2a, Suppl. fig. 3–4). HybridCheck further identified an EAV-like segment present in PRRSV (Suppl. fig. 5). Signal patterns and quartet configurations strongly suggest exchange with an unsampled arterivirus closely related to EAV, rather than direct equine-to-swine transmission, and exclude reverse transfer from PRRSV into EAV.

**Figure 2.**
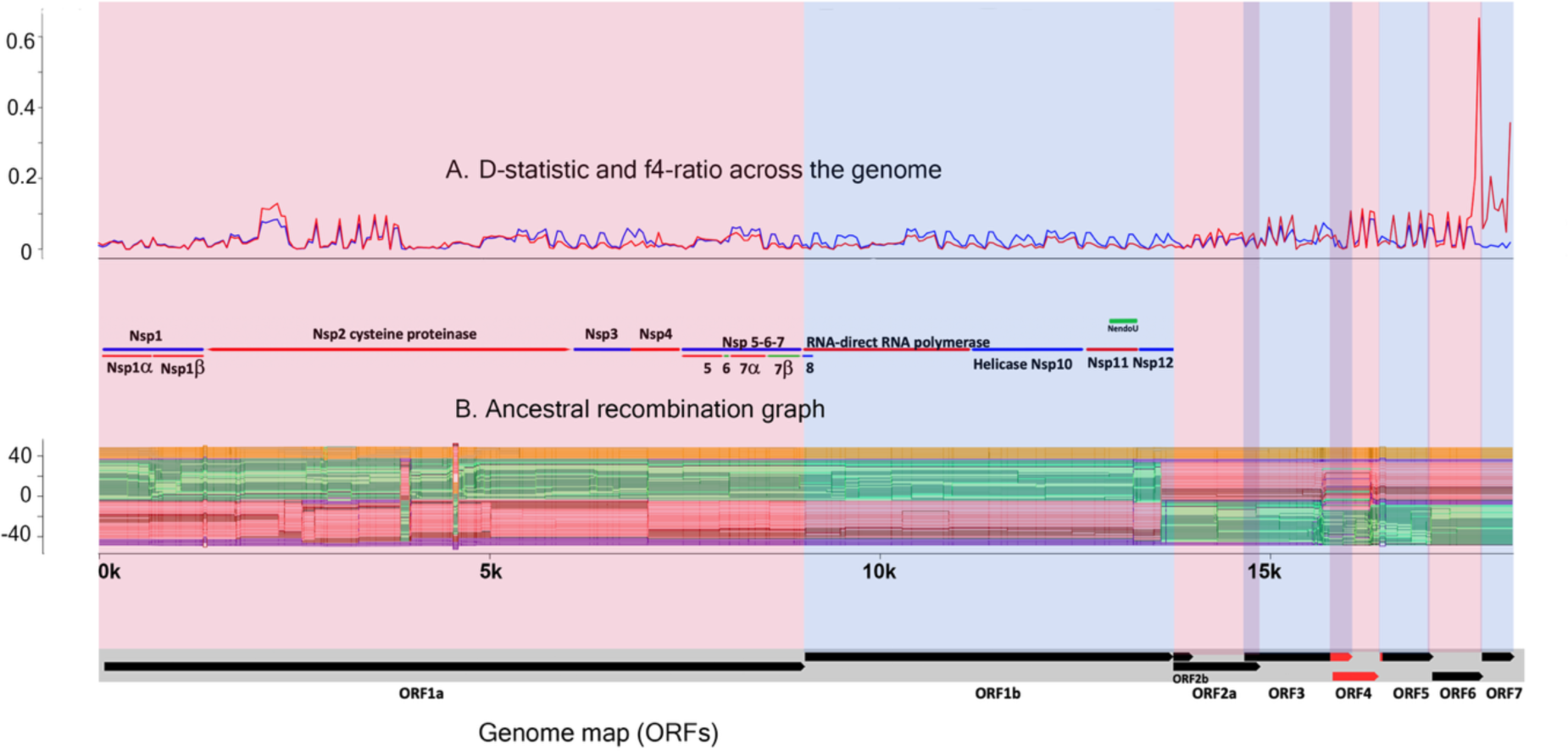
Recombination and introgression signals. A) D-statistic and f4-ratio across the genome; B) Ancestral recombination graph.

### Recombination and cross-species introgression shape the evolutionary mosaic of PRRSV

#### Genome-wide recombinations signal

Genome-wide analyses revealed pervasive mosaicism across PRRSV genomes, with recombination signals detected in most coding regions. To minimise false positives, we adopted a conservative consensus framework: only breakpoints supported by at least three independent algorithms, corrected for multiple testing, and corroborated by phylogenetic incongruence were retained. While GARD, which infers recombination from shifts in phylogenetic likelihood across sequence alignments, did not identify statistically significant breakpoints (Suppl. fig), 3SEQ, a method that detects mosaic structures by testing all possible triplets of sequences, recovered numerous events spanning nearly the entire genome. RDP5, which applies a suite of complementary algorithms within a single framework, pinpointed a robust signal in ORF4 linking representative general European and Danish isolates (Suppl. fig. 6).

#### Hotspot versus conserved regions

Recombination-aware inference with ARGweaver, which reconstructs local genealogies to detect ancestry switching, revealed a heterogeneous landscape of alternating conserved blocks and highly mosaic segments. The strongest and most consistent signals localise to ORF4 (GP4) (Fig. 2b), which displays unusually deep local TMRCA and multiple parental contributions, consistent with repeated domain swapping (Suppl. Fig. 6). Additional recombination segments were detected in ORF1a and ORF5 (GP5). In contrast, ORF1b, together with the structural genes ORF6 (M) and ORF7 (N), showed little to no evidence of recombination, probably reflecting stronger functional constraints (Fig. 2a–b, Suppl. Fig. 3–5). Weak signals in ORF1b detected by other methods were not supported by ARGweaver or breakpoint analyses and are therefore treated as non-decisive. Complementary Patterson’s D and f₄-ratio tests (ABBA–BABA framework), which identify excess shared nucleotide patterns as evidence of introgression beyond incomplete lineage sorting, confirmed significant gene flow between PRRSV and rodent arteriviruses, with reproducible signals in ORF1a, ORF4, and ORF5. Together, these results identify GP4 and GP5 as recurrent hotspots of recombination and introgression, while polymerase and internal structural genes remain comparatively less prone to recombination, or at least to recombination that has been fixed in the genome.

#### Cross-species introgression

Species-tree reconciliation with GeneRax, which detects incongruences between gene trees and species trees as signatures of horizontal transfer, revealed two main patterns: (i) ancient introgression from rodent arteriviruses into the ancestors of both PRRSV-1 and PRRSV-2 (Suppl. Fig. 7), and (ii) more recent exchanges linking extant PRRSV clades with rodent viruses (Suppl. Fig. 7). These signals are concentrated in ORF4, with secondary peaks in ORF1a and ORF5 (Fig. 2b; Suppl. Fig. 8). Complementary HybridCheck analyses detected an EAV-like segment in both PRRSV species, though at distinct coordinates in ORF1a for PRRSV-1 and PRRSV-2 isolates. Quartet configurations and directionality tests indicate that this tract most likely originated from an unsampled arterivirus closely related to EAV, rather than from modern equine arteriviruses, and exclude reverse transfer into EAV. Together, these results extend the recombination landscape beyond within-lineage processes, highlighting repeated cross-species introgression as a key contributor to the mosaic architecture of PRRSV genomes.

#### Lineage anomalies and the mosaic architecture of PRRSV

Among the clearest illustrations of PRRSV’s mosaic evolution is an early European PRRSV-1 lineage (Olot-91; Spain, 1991 ^23^), which carries multiple recombination breakpoints despite its putative parental sequences being sampled at later dates. This anomaly is most parsimoniously explained by recombination with unsampled or extinct intermediates, underscoring how incomplete sampling can distort apparent genealogical relationships. Comparable patterns occur sporadically across other isolates, suggesting that many observed recombinants represent only fragments of a broader, unsampled network of ancestral diversity.

Taken together, our analyses show that recombination and cross-species introgression are central forces in PRRSV evolution (Fig 2a-b), producing pervasive genomic mosaics that cannot be captured by simple tree-like models (Suppl.fig 7-8). The envelope glycoproteins GP4 and GP5 consistently emerge as hotspots, in line with their roles in receptor binding and immune evasion, whereas polymerase and structural genes M and N remain comparatively recombination-poor. Beyond within-lineage exchanges, recurrent introgression from rodent arteriviruses and an EAV-like source has contributed to both PRRSV species. Collectively, these findings establish PRRSV as a virus shaped not by a single trajectory of divergence, but by a reticulate evolutionary history driven by recombination, introgression.

### Contrasting selective pressures across the PRRSV genome

Codon-based analyses reveal a highly uneven distribution of selective pressures across the PRRSV genome, with surface glycoproteins showing recurrent adaptive diversification, whereas *replicase*-associated and structural proteins M and N remain dominated by purifying constraint (Fig. 3a-b). Pervasive codon-level tests (FEL, SLAC, FUBAR, which estimate whether codons are under constant selection across the tree) (Suppl.fig 9) consistently identify ORF3 as the principal focus of positive selection, with secondary signals in ORF2a and occasional sites in ORF7 (N). In contrast, ORF6 (M) shows no evidence of diversifying selection and ORF1b contributes only a few positively selected sites. Most codons genome-wide fall under negative selection, with the lowest proportions of constrained sites in ORF3 and ORF2a, underscoring their relative lability.

**Figure 3.**
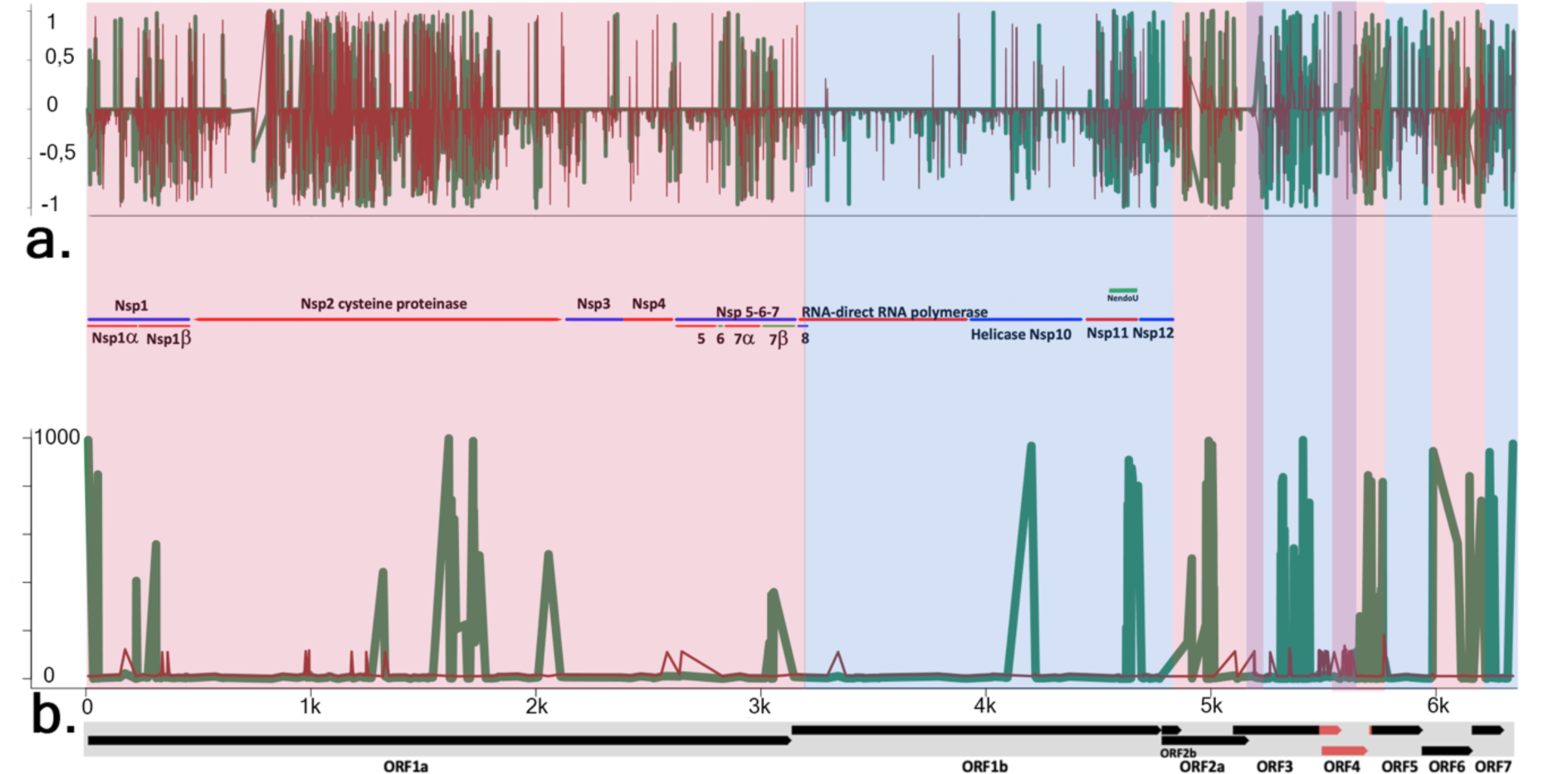
Selection pressures across the PRRSV genome. A) Site-specific selection inferred using FEL. Positive values indicate diversifying (positive) selection, while negative values indicate purifying (negative) selection. B) Episodic selection inferred using MEME. Peaks indicate codons experiencing episodic diversifying selection. Gene boundaries and key non-structural proteins (nsp) are shown below for reference, The green line represents results obtained for the entire *Arteriviridae* clade, whereas the thin red line corresponds to analyses restricted to PRRSV sequences.

Method concordance was high. SLAC and FUBAR broadly agreed with FEL, while episodic tests (MEME, which detects bursts of selection at individual codons, and aBSREL/BUSTED, which identify branches or genes experiencing episodic diversification), highlighted adaptive episodes in ORF2a, ORF3, and ORF5. This is consistent with immune-driven diversification. In contrast, ORF7 (N) shows sporadic positively selected sites depending on method, which might reflect primarily structural constraints with occasional adaptive episodes.

Comparisons between clades point to species-specific behaviour. Relative to other *arteriviruses*, the PRRSV clade exhibits stronger purifying selection in ORF4 and ORF7, consistent with ORF4’s oligomeric scaffold function and the structural constraints on N protein. ORF3 contains fewer neutrally evolving sites yet undergoes frequent episodes of both positive and negative selection. Between species, PRRSV-1 shows relaxed constraint in ORFs 5–7, whereas PRRSV-2 displays intensified diversifying selection in ORF5 but stronger conservation in replicase genes. Episodic branch-site analyses (aBSREL) further identify intensified selection in ORF3 and ORF5 along branches leading to highly virulent Chinese isolates, while BUSTED supports gene-wide diversification in these loci. REL-family analyses (which model variation in selective pressure across sites and clades) highlight clade-specific intensification in ORF5, consistent with its role as a major neutralizing epitope (Suppl.fig 9)

Together, these results establish ORF3 and ORF5 as recurrent hotspots of adaptive diversification, driven by immune escape, while ORF2a and ORF7 provide secondary but consistent signals. By contrast, ORF1a/1b, ORF6, and much of ORF4 remain under strong purifying constraint, forming a conserved genomic backbone. The contrasting signatures between PRRSV-1 and PRRSV-2 suggest that species-specific immune landscapes and structural constraints shape distinct adaptive trajectories.

Genome-wide codon-based analyses reveal strong positive selection in ORF3, while replicase-associated genes (ORF1a/1b) remain highly conserved. Purifying selection dominates in ORF6 and ORF7, consistent with their structural roles. These patterns highlight ORF3 as the main target of adaptive diversification.

### Human-mediated demographic expansion of PRRSV

#### Viral demographic reconstructions

Bayesian coalescent models (skyline, skyride, skygrid) reveal that PRRSV demographic history closely parallels changes in pig production (Fig 4, Suppl.fig 10-11). The two lineages show distinct trajectories: In contrast to rodent *arteriviruses*, which remained demographically stable, PRRSV-1 expanded sharply around the 1989 Western European outbreak, while PRRSV-2 rose during the 1983 wave of outbreaks across US Midwest and later declined. Excluding recombinant and positively selected regions produced smoother yet consistent curves, which confirmed lineage-specific expansions. ARG-based reconstructions add a genomic dimension, revealing higher local effective population sizes (Ne) and deeper genealogical depths in ORF1a/1b. In contrast, ORF3, ORF5, and ORF7 display shallow TMRCAs, reflecting purifying selection and more recent divergence.

**Figure 4.**
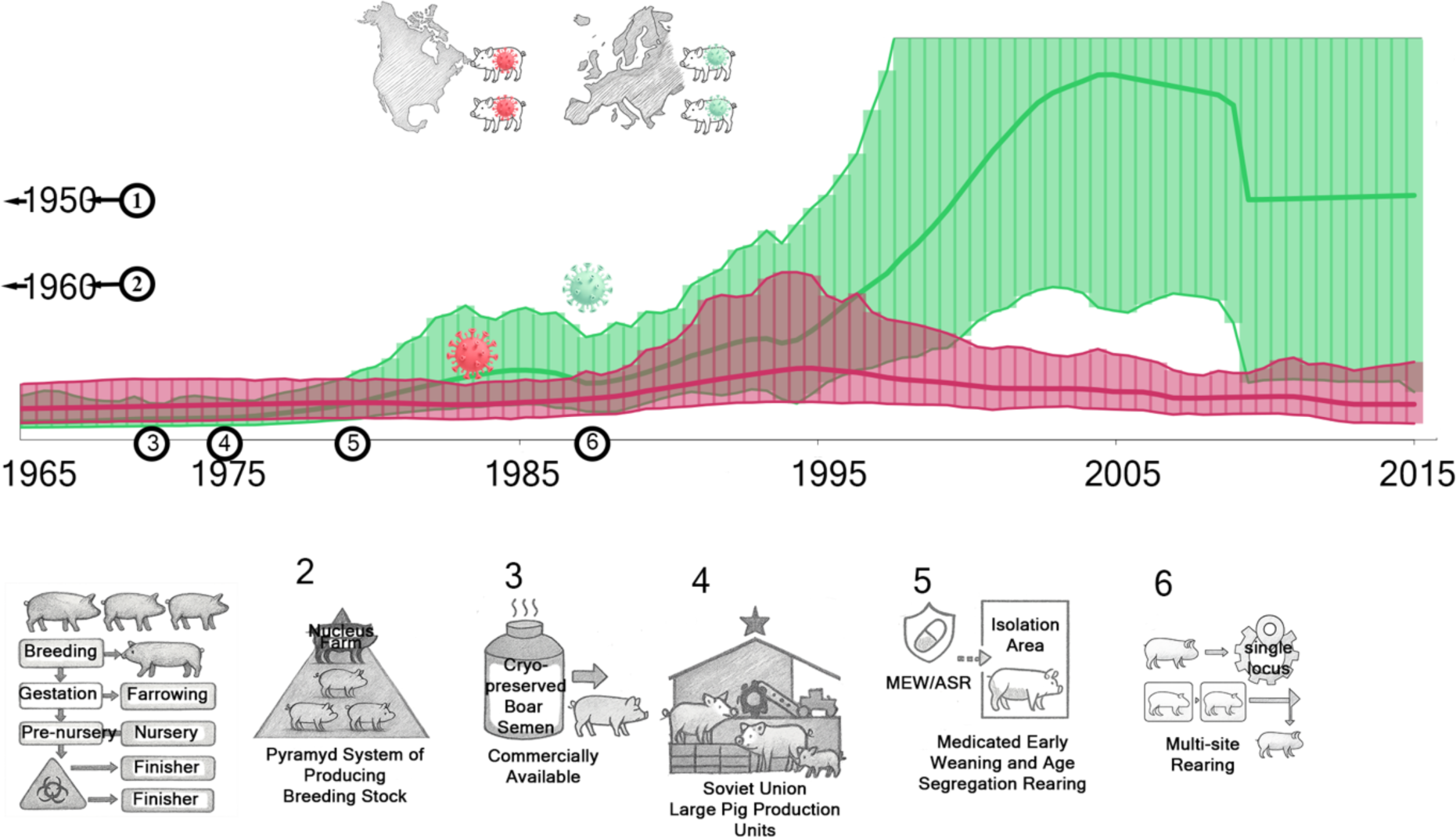
Demographic history and human-mediated drivers of PRRSV evolution. Bayesian skyline plots show population expansions of PRRSV-1 and PRRSV-2 coinciding with mid-20th century intensification of pig farming. Pig production statistics confirm rapid increases in herd size despite declining breeding populations. Conceptual summary (right) illustrates rodent origins of PRRSV, subsequent divergence of PRRSV-1 and PRRSV-2 lineages, and human-driven amplification of viral diversity. Green Skyplots is PRRSV-1 wile red is PRRSV-2.

#### Parallel host demography

These viral dynamics unfolded against a backdrop of rapid changes in pig production (Suppl.fig 12). Between 1965 and 2000, total pig numbers in the U.S. and Western Europe rose even as the number of breeding herds decreased (Suppl.fig 13), driven by sustained productivity gains per sow (Suppl.fig 13). Regional contrasts were pronounced: the number of pigs plateaued in Germany, Denmark, and the Netherlands after the late 1980s, while Spain continued to expand. Density maps show that early outbreak zones the U.S. Midwest and Lower Saxony were among the densest pig-farming regions on their respective continents (Suppl.fig 14).

#### Husbandry innovations and viral amplification

Historical innovations in husbandry further aligned with these demographic shifts. Confined rearing in the 1950s, pyramid breeding systems in the 1960s–70s, and the adoption of artificial insemination and cryopreserved semen by the late 1970s accelerated herd turnover and connectivity. By the 1980s, medicated early weaning and multi-site rearing fragmented age cohorts and scaled up production, while large, mechanised systems came to dominate both continents. Together, these changes created dense, highly connected host populations that amplified viral spread, imposed transmission bottlenecks, and facilitated the fixation of adaptive variants. The convergence of viral recombination, selection, and mid-20th-century farming innovations transformed an arterivirus of uncertain ancestry – most closely related to rodent lineages – into one of the most economically damaging pathogens of modern swine production.

**Figure 5.**
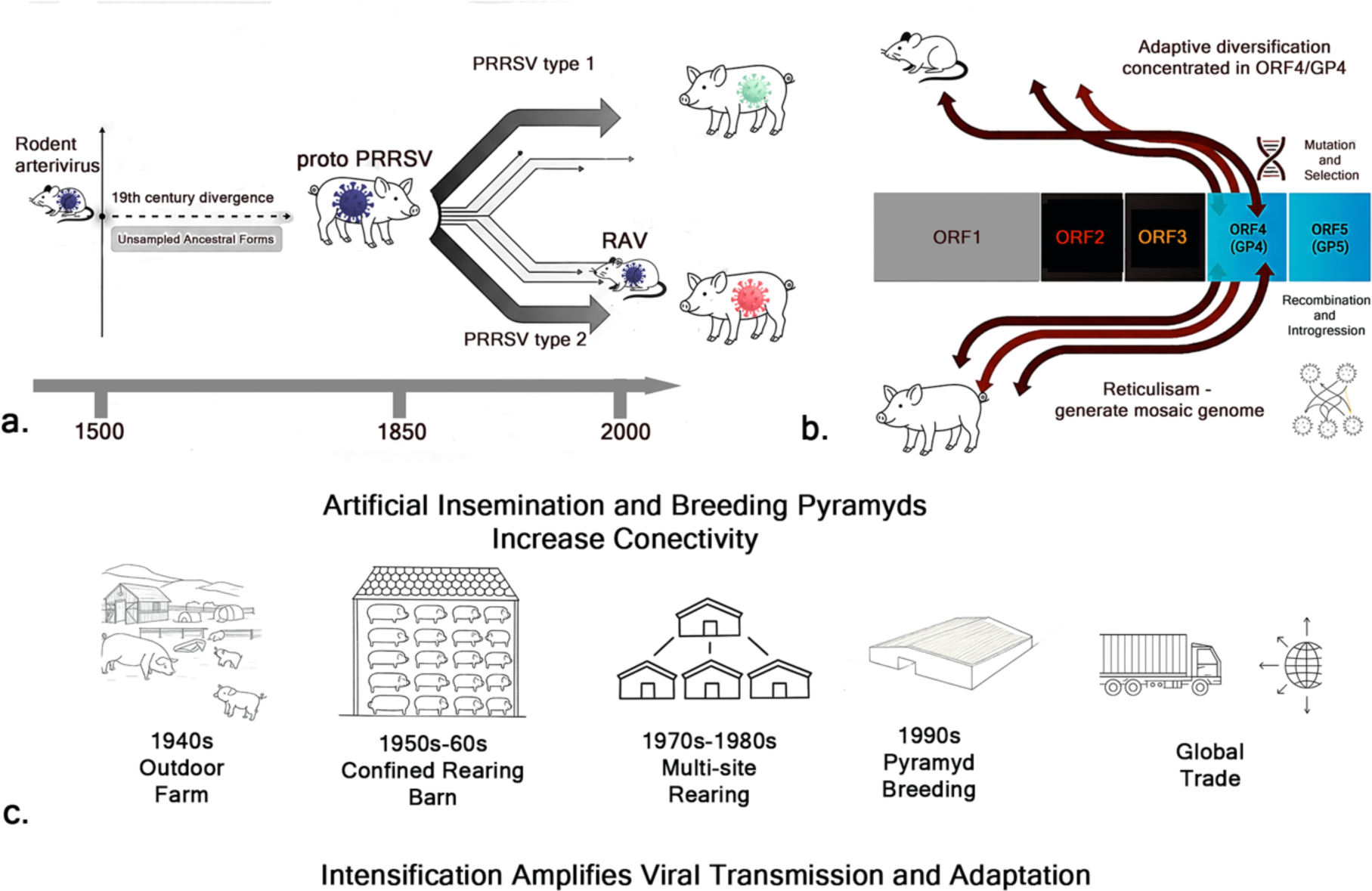
Integrative model of PRRSV evolution and emergence. Schematic summary illustrating how genomic, ecological and anthropogenic processes interacted to shape the evolutionary history of PRRSV. Viral icons denote major lineages: red indicates PRRSV-2, green PRRSV-1 and purple rodent arteriviruses. A) Evolutionary origins. A Bayesian time-calibrated reconstruction places the divergence between PRRSV-1 and PRRSV-2 lineages in the early 19th century, well before recognized outbreaks. This split occurs after the separation of rodent arteriviruses, with unsampled ancestral forms likely bridging the gap between the rodent and proto-PRRSV lineages. B) Genomic mechanisms. Recombination, introgression and positive selection generate a mosaic genomic architecture, with adaptive diversification concentrated in the envelope glycoproteins GP4 and GP5. These loci act as recurrent hubs for genetic exchange, whereas replicase genes remain comparatively conserved. C) Anthropogenic drivers. Mid-20th-century intensification of pig production, characterized by confined rearing, artificial insemination, multi-site systems, breeding pyramids and global trade, transformed pig populations into dense, genetically interconnected metapopulations. This restructuring amplified viral transmission, facilitated the fixation of adaptive variants and enabled the global persistence of PRRSV.

## Discussion

The origins of PRRSV, and the circumstances that led to the emergence of two closely related species almost simultaneously on different continents, have long been subjects of debate and controversy. Emerging viruses often arise through the interplay of wildlife reservoirs, genomic innovation, and human-driven ecological change. Using an integrative phylogenomic framework that jointly considers recombination, selection, and demography, we show that the European (PRRSV-1) and North American (PRRSV-2) lineages diverged in the early nineteenth century, long before the disease was recognized, and that their closest extant relatives are rodent-associated arteriviruses, consistent with a prolonged pre-epidemic phase. Recombination and introgression have produced a mosaic genome and a reticulate evolutionary history, with adaptive diversification concentrated in the surface glycoproteins GP4, GP5, and ORF3, while the replicase complex forms a conserved genomic backbone. Demographic reconstructions further link the mid-twentieth-century intensification of pig production to viral population expansion. Together, these results position PRRSV as a model for how evolutionary processes and human practices converge to drive viral emergence within intensive livestock systems.

Early molecular clock studies based on single-gene phylogenies dated the split between PRRSV-1 and PRRSV-2 to 1979–1980 ^17,24^, consistent with the description of the first recognized PRRS outbreaks. Forsberg (2005) initially obtained an implausibly early divergence estimate around 1593, but this reflected the use of a limited genomic region strongly affected by selection and recombination. When he later focused on an expanded ORF3 dataset from an appropriate temporal window, avoiding biases due to recombination and selection, he arrived at a more credible estimate for the divergence between the two species that was dated back to ∼1880 ± 15 years ^18^, indicating pre-epidemic separation. Our whole-genome analysis, which explicitly excluded all sites showing signatures of positive selection or recombination, independently converges on the same late 19th-century divergence time but with smaller confidence intervals (± 6 years). This concordance underscores how methodological choices - especially the genomic regions analysed - critically influence molecular clock estimates ^25,26^. Widely used loci such as ORF5 and ORF7, while valuable for diagnostics, are problematic for phylogenetic inference: ORF5 is shaped by selection and recombination, whereas ORF7 is only constrained by selection. Thus, any reliance on these loci would contribute to inconsistent divergence estimates. Collectively, these findings strongly support the existence of a pre-epidemic reservoir, either in another species or in pigs long before recognized disease emergence, from which multiple outbreaks likely occurred ^17^.

Although PRRSV was initially thought to have originated in North America and then spread to Europe, phylogenetic diversity patterns point to Eastern Europe as the likely source of the European epidemic ^16^. Stadejek et al., ^16^ proposed that geopolitical changes following the fall of the Berlin Wall facilitated westward viral spread, aligning viral movement with historical trade and livestock flows across a newly opened Europe. A critical new piece of evidence came with the identification of a rodent *arterivirus* isolated from rats in China in 2016 ^27^, which exhibits a closer phylogenetic relationship to North American PRRSV strains than to European ones ^22^. This discovery supports the existence of a broader rodent - PRRSV radiation before the first recognized PRRS epidemics and implicates rodents as long-term viral reservoirs and genetic exchange partners, rather than transient intermediates. Contrary to earlier proposals linking the divergence of PRRSV-1 and PRRSV-2 to the transatlantic movement of pigs ^28^, our analyses find no phylogenetic support for such a scenario. Instead, the deep split may more likely reflect the separation of a proto-PRRSV circulating in pigs (derived from its rodent relatives) during the formation and global dissemination of modern pig breeds in the 18th–19th centuries. The establishment and export of standardized breeds, together with early livestock exhibitions ^29^, provided ideal conditions for the silent spread of a subclinical virus. Our speculation is that the newly discovered Chinese rodent *arterivirus* is best interpreted as a closely related lineage involved in recurrent genetic exchange, rather than a direct ancestor. Likewise, the hypothesis of westward PRRSV spread within Europe after the fall of the Berlin Wall is less plausible than earlier east-to-west or Europe-to-North America dissemination, consistent with higher genetic diversity in European lineages. Eastern Europe, where large state/collective pig-production systems were established earlier (1920s–1930s), may have functioned as an early amplification zone, consistent with higher within-farm densities, reduced herd segmentation and increased movement between integrated units ^30^. Overall, PRRSV diversification seems to be unfolded during a prolonged pre-epidemic phase, whereas clinical emergence was probably triggered later by husbandry shifts, not sudden evolutionary leaps.

Demographic reconstructions indicate that the mid-20th century intensification of pig husbandry imposed repeated bottlenecks, amplifying genetic drift and accelerating the fixation of advantageous variants. Population bottlenecks can paradoxically accelerate viral adaptation: when few genomes seed subsequent generations, drift can overpower selection, allowing rare beneficial variants to “surf” to high frequency before sweeping to fixation once populations rebound. Such bottleneck-driven accelerations have been demonstrated in multiple RNA viruses ^31,32^. In PRRSV, repeated transmission bottlenecks during herd expansion likely purged standing diversity while amplifying adaptive mutations, setting the stage for rapid post-bottleneck rebounds. Unusually, both PRRSV-1 and PRRSV-2 lineages exhibit clear bottleneck signatures coinciding with their respective outbreak periods. Available evidence suggests that the virus circulated cryptically before disease recognition, but post-bottleneck expansions acted as evolutionary accelerants, aggregating rare or structurally constrained variants and triggering epidemic emergence. Thus, the European and North American outbreaks represent parallel evolutionary responses to analogous demographic triggers rather than independent evolutionary leaps.

Recombination and introgression emerge as central forces in arterivirus evolution, generating pervasive reticulation that challenges simple tree-like models of diversification. Comparable processes are seen across RNA viruses, including coronaviruses and influenza viruses, where segmental exchange or recombination repeatedly reshapes genomes and drives antigenic innovation ^26^. In PRRSV, the resulting genomic reticulism, most pronounced in GP4 and GP5, illustrates how structural and immune-exposed glycoproteins act as recurrent hubs for genetic exchange. Variation in receptor-binding domains and neutralizing epitopes provides a direct mechanistic link between recombination, host adaptation, and immune escape. In contrast, the replicase complex and internal structural proteins remain under strong purifying constraint, forming a conserved genomic scaffold that anchors this flexibility. This asymmetric genome architecture, adaptive envelope genes embedded within a stable replication core, appears to represent a recurring strategy by which arteriviruses balance evolutionary innovation with functional integrity.

Yet this intrinsic genomic plasticity alone cannot explain the success and global spread of PRRSV. Our demographic reconstructions reveal that its expansion closely paralleled the industrialization of pig production in the mid-twentieth century, when confined rearing, artificial insemination, and international breeding networks transformed swine populations into dense, genetically interconnected metapopulations. These human innovations amplified transmission opportunities, shortened generation intervals, and intensified immune selection. At the same time, strong artificial selection for production traits eroded immune gene diversity, homogenizing host defences and narrowing the adaptive landscape available to the virus.

Despite the integrative scope of our analyses, several limitations remain. Evolutionary reconstructions depend on available genomes, whose temporal and taxonomic gaps may bias divergence estimates and obscure unsampled intermediates. Access to historical viral material could clarify early diversification, but such samples are scarce, and the RNA nature of PRRSV further complicates recovery. ARG-based inference, while powerful, remains computationally intensive and alignment-sensitive. Continuous NGS-based surveillance, systematic temporal sampling, and integration of host genomic data will be essential to refine evolutionary timelines, detect emerging recombinants, and illuminate host–virus coevolution. Ultimately, this study demonstrates the value of a systems-level, One Health perspective for understanding viral emergence. By coupling phylogenomic with demographic and anthropogenic context, we reveal how genomic, ecological, and human processes intersect across biological scales. This framework is broadly applicable to other rapidly evolving RNA viruses and provides a basis for predictive, genome-informed surveillance and control strategies at the human–animal interface.

## Materials and Methods

### Genome data and taxon sampling

To survey the broader diversity of arteriviruses with a focus on PRRSV, we compiled 1,818 complete genome sequences from GenBank (accessed 15 October 2025). The dataset comprised: 95 equine arterivirus (EAV), 41 simian haemorrhagic fever virus and related simian arteriviruses (SHFV), 28 lactate dehydrogenase–elevating virus and related rodent arteriviruses (LDEV), 2 forest pouched giant rat arterivirus (APRAV), 4 rat arterivirus (RAV), 189 PRRSV type 1, and 1,459 PRRSV type 2 genomes.

### Alignment and phylogenetic reconstruction

Complete genomes were aligned with MAFFT. A Neighbour-Joining phylogeny was inferred in MEGA, and the tree was visualized and annotated in iTOL. Where relevant, poorly aligned regions were inspected and masked prior to tree inference.

### Time-scaled analysis (deep evolution of PRRSV)

To investigate the deeper evolutionary history of PRRSV, we selected a representative subset of 114 complete genomes that included major PRRSV lineages and appropriate outgroups from other arteriviruses. Using this subset, we inferred a time-scaled phylogeny to estimate divergence times among principal PRRSV clades, ensuring adequate sampling across lineages and minimizing sampling-date/sequence-length confounding. Taxon names for non-PRRSV sequences followed the format “Virus/Gene accession number_year of isolation.” For PRRSV sequences, additional information on the country of origin was included in the format “PRRSVtype/Country of origin_Gene accession number_year of isolation.” Reference strains Lelystad virus and VR2332 were specifically labelled. Complete genome sequences were aligned using ClustalW in MEGA 6 with a codon-based alignment. Sequences were translated into amino acids, re-aligned, and subsequently back translated to nucleotides to facilitate protein identification and check for sequencing errors. The complete genome alignments were partitioned into individual ORFs (ORF1a–ORF7). Each ORF was extracted from the full genome alignment, re-aligned, translated into amino acids, and processed to remove all stop codons as potential sequencing errors ^33^. Complete genome and partitioned alignments were exported in .nexus format.

### Recombination analysis

Recombination events were identified using multiple complementary methods. We first applied the Single Breakpoint Recombination (SBP) and Genetic Algorithm for Recombination Detection (GARD), both implemented in HyPhy ^34^, under a General Time Reversible (GTR) model with gamma-distributed rate heterogeneity. Analyses were run with 100 bootstrap replicates, and breakpoints were retained if the no-recombination model was rejected by a decrease in AIC greater than 100. The same alignments were then analysed with the 3SEQ recombination detection algorithm ^35^, executed via the Unix command-line on Linux and macOS in “full run” mode with a 700 × 700 × 700 p-value table. Putative recombinant fragments were retained only if they showed mosaic signals with a Dunn–Šidák–corrected P < 0.01, and their parental sequences were recorded. Finally, the dataset was analysed in RDP5 ^36^, which integrates seven different algorithms (RDP, GENECONV, MaxChi, BootScan, SiScan, Chimaera and 3SEQ) in a Windows graphical interface. Analyses were performed under the highest-stringency settings with Bonferroni-corrected P < 0.05. A recombination event was accepted as high confidence only when three criteria were met: (i) its breakpoints lay within 200 nucleotides of a GARD-predicted site; (ii) the same fragment was significant in 3SEQ; and (iii) it was independently detected by at least three of the RDP5 algorithms with consistent parental assignment.

### Selection Pressure Analysis

Selection pressure was analysed for both the complete arterivirus dataset (AV) and a subset comprising PRRSV and RAV sequences (PRRSV). Multiple complementary methods were applied to evaluate evolutionary dynamics across open reading frames (ORFs).

Codon-level selection was first screened using a two-step workflow. In the initial rapid scan, FEL (Fixed Effects Likelihood, P < 0.05) and SLAC (Single-Likelihood Ancestor Counting, 100 ancestral reconstructions, P < 0.10) ^37^ were used to identify candidate codons. Each candidate site was then re-examined with the more specific FUBAR (Fast, Unconstrained Bayesian Approximation, five Metropolis–Hastings chains, 2 × 10⁶ iterations, Dirichlet prior = 0.5) to detect pervasive selection across the tree, and MEME (Mixed Effects Model of Evolution, P < 0.01) ^38^ to capture episodic positive selection on individual branches. A codon was accepted as positively selected only if it was first identified by FEL or SLAC and subsequently confirmed by either FUBAR or MEME. This combined strategy balanced the sensitivity of the initial scan with the specificity of the confirmatory tests, reducing false positives while retaining weak or lineage-specific signals.

To assess selection dynamics above the codon scale, three branch- and clade-level tests were applied. RELAX was used to evaluate whether the overall strength of selection had weakened (indicating possible functional decay) or intensified (suggesting adaptive optimisation). To distinguish between adaptive change and increased purifying pressure, branches showing intensification were further examined with aBSREL (adaptive Branch-Site Random Effects Likelihood, three ω-rate classes) ^39^ and BUSTED (Branch-Site Unrestricted Statistical Test for Episodic Diversification) ^36^. Only when aBSREL detected episodic ω > 1 or BUSTED returned a significant clade-wide signal (FDR-adjusted P < 0.05) was the shift classified as an adaptive trend; otherwise, it was recorded as heightened purifying selection.

Together, this multi-layered pipeline combined the global sensitivity of RELAX with the specificity of aBSREL and BUSTED, allowing us to chart both functional loss and lineage-specific adaptation across the arterivirus phylogeny.

### Bayesian phylogeny inference

Bayesian phylogenies were inferred in BEAST v1.10.4 ^40^, with Markov Chain Monte Carlo (MCMC) chains run until effective sample sizes (ESS) exceeded 200. Alternative substitution models (GTR+Γ4+I, HKY+Γ4+I, and codon models SRD06 and YANG96), clock schemes (strict and uncorrelated relaxed clocks with lognormal, gamma, or exponential distributions, as well as random and fixed local clocks), and demographic priors (constant, exponential, expansion, logistic, and Bayesian skyline) were compared. Model fit was evaluated using the Akaike’s Information Criterion for MCMC samples (AICM) implemented in *Tracer* v1.6 ^41^. The optimal combination of models was then re-run on alignments filtered to remove recombinant and positively selected sites. Posterior distributions were summarised as maximum clade credibility (MCC) trees, while neighbour-joining (NJ) trees were generated in parallel to validate overall topologies.

### Phylogenetic trees

A range of phylogenetic scenarios were explored to assess model choice, the effect of alignment partitioning, and the impact of recombination or selection on tree topology. These included NJ trees on the full dataset, MCC trees inferred under both nucleotide and codon substitution models, and trees based on individual ORFs as well as filtered genome partitions. Pre-analysis screening identified ORF6, encoding the major envelope protein, as the best compromise between phylogenetic depth and clock-like behaviour, and therefore it was used as the primary focus for time-calibrated analyses. The full set of alternative trees is presented in the Supplementary Information (Suppl.fig 2, while the representative MCC tree from ORF6 is shown in Suppl.fig 1.

### Introgression analyses

Introgression was assessed using three independent methods. *HybridCheck* ^42^, implemented in R, was used for rapid detection, visualisation and approximate dating of recombinant regions. *Dsuite* ^43^ was employed to calculate Patterson’s D-statistic (also known as the ABBA–BABA test) and related metrics such as the f₄-ratio, which are widely used to detect gene flow between populations or closely related species. Owing to its computational efficiency, Dsuite enabled genome-wide evaluation of all trio combinations across the dataset. Finally, *ARGweaver-D* ^44^ was used to reconstruct the ancestral recombination graph (ARG), a framework regarded as the most complete representation of reticulate evolutionary history ^45^. Together, these methods allowed us to capture both recent mosaic tracts and deeper, lineage-spanning introgression events.

### HybridCheck Analysis

*HybridCheck* was applied to test predefined taxon triplets for mosaic genome blocks. Eight representative triplets spanning rodent, simian and porcine *arteriviruses* were analysed with appropriate outgroups, allowing us to visualise and date potential introgression events across deep lineages. Analyses were run with both broad and fine-scale scanning windows. Full details of the tested triplets, abbreviations, parameter settings and input files are provided in the GitHub.

### Dsuite analysis

*Dsuite* was used to quantify introgression signals across the arterivirus phylogeny. Divergent primate viruses were excluded as unlikely to provide meaningful comparisons with porcine lineages. Genome-wide Patterson’s D-statistics, Z-scores, p-values and f₄-ratios were computed with *Dtrios*. Localised introgression was scanned with *Dinvestigate* (200 nt window, 25 nt step), while *Fbranch* was used to map clade-specific f₄-ratio signals onto the phylogeny. Based on prior phylogenetic analysis, PRRSV-1 was subdivided into three clades, PRRSV-2 into four, and EAV into four, providing a framework for lineage-level tests. Full details of the excluded taxa, clade definitions and input files are provided in the GitHub.

### ARGweaver

The ancestral recombination graph (ARG) was reconstructed with *ARGweaver-D* using the same sequence set as in the Dsuite analysis. Parameters followed Bayesian point estimates: mutation rate = 2.3 × 10⁻³ substitutions per site per year, recombination rate = 1.3 × 10⁻³ to 1.3 × 10⁻¹² events per site per year, and effective population size ranging from 7.57 × 10⁰ to 10⁶. Markov Chain Monte Carlo (MCMC) analyses were run for 20,000 iterations until convergence. Sequences were treated as haploid FASTA files and tip-dated using an age file. The results were visualised in R, with clades colour-coded to match the Bayesian phylogeny for clarity of introgression blocks. Input FASTAs, age files and command lines are provided in the GitHub.

#### GeneRax reconciliation

Species-aware gene tree reconciliation was performed with *GeneRax* ^46^, using alignments partitioned according to recombination blocks inferred by *ARGweaver-D* as well as full ORFs. In the block-level analysis, the genome was divided into 26 segments spanning ORF1a through ORF7, each treated as an individual gene. In the ORF-level analysis, each complete ORF was analysed as a single gene. For both datasets, maximum likelihood gene trees were reconciled against the species tree inferred from the Bayesian analysis. Reconciliations were visualised with *ThirdKind* ^47^, which generates scalable vector graphics indicating transfers, duplications and losses. All partition definitions, input FASTAs and command lines are provided in the GitHub.

#### Population dynamic

Trends in pig production were obtained from EUROSTAT (EU-27) and USDA (1965–2020) databases. Viral demographic history was reconstructed from a PRRSV dataset comprising 37 European (PRRSV-1) and 38 North American (PRRSV-2) genomes.

### Skyplots

Viral demography was inferred using three coalescent-based models implemented in *BEAST*: Bayesian Skyline, Bayesian Skyride and Bayesian SkyGrid. For calibration, the mean tree height of the PRRSV dataset was set as the node height of the PRRSV branch, with the standard deviation adjusted to encompass the 95% highest posterior density (HPD). Analyses were run on two datasets: (i) the complete PRRSV dataset including PRRSV-EU, PRRSV-US and RAV sequences; and (ii) a filtered subset with codons flagged as recombinant (3SEQ) or under selection (FEL) removed.

## Data availability

All sequence data analysed in this manuscript are available at https://github.com/dinkonovosel/Supplementary-information-Rodent-Origins-and-Human-Mediated-Evolutionary-Dynamics-of-PRRSV.

## Code availability

All custom code used in the manuscript is available at https://github.com/dinkonovosel/Supplementary-information-Rodent-Origins-and-Human-Mediated-Evolutionary-Dynamics-of-PRRS.

## Acknowledgments

The contribution of Dinko Novosel and Ino Curik to this work was supported by the Croatian Science Foundation under project number HRZZ IP-2022-10-8926. We are grateful to Silvana Čubrić for her creative support and expert assistance in the graphical design and refinement of the figures.

## Author contributions

All authors contributed to analyses and interpretations. D.N performed recombination, introgression, demographic analysis. D.N. and O.R.S. performed phylogenetic analysis. D.N., I.C. and A.J. prepared graphic illustrations. D.N. wrote the first draft of the manuscript, and all authors contributed to manuscript editing. I.C. and E.M. jointly supervised the study and share senior authorship.

## Competing interests

The authors declare no competing interest

## Notes

https://github.com/dinkonovosel/Supplementary-information-Rodent-Origins-and-Human-Mediated-Evolutionary-Dynamics-of-PRRS

### Competing Interest Statement

The authors have declared no competing interest.

## References

1. Loula, T. Mystery pig disease. Agri-practice 23–34 (1991).

2. Keffaber, K. K. Reporductive failure of unknown etiology. Am. Assoc. Swine Pr. Newsletters 1–10 (1999).

3. Lunney, J. K., Benfield, D. A. & Rowland, R. R. R. Porcine reproductive and respiratory syndrome virus: an update on an emerging and re-emerging viral disease of swine. Virus research 154, 1–6 (2010).

4. Holtkamp, D. J. et al. Assessment of the economic impact of porcine reproductive and respiratory syndrome virus on United States pork producers. Swine Heal. Prod. 21, 72–84 (2013).

5. Busse, F. W., Alt, M., Janthur, W., Neuman, P. & Schoss, P. Epidemiological studies on porcine epidemic abortion and respiratory syndrome (PEARS) in Lower Saxony of Germany. in Proceedings of 12th Congress International Pig Veterinary Society, The Netherlands 115 (1992).

6. Wensvoort, G. et al. Mystery swine disease in The Netherlands: the isolation of Lelystad virus. Vet. Q. 13, 121–130 (1991).

7. Collins, J. E. et al. Isolation of swine infertility and respiratory syndrome virus (isolate ATCC VR-2332) in North America and experimental reproduction of the disease in gnotobiotic pigs. J. Vet. Diagn. Invest. 4, 117–126 (1992).

8. Snijder, E. J. The arterivirus replicase. The road from RNA to protein(s), and back again. Adv. Exp. Med. Biol. 440, 97–108 (1998).

9. Wu, W.-H. et al. A 10-kDa Structural Protein of Porcine Reproductive and Respiratory Syndrome Virus Encoded by ORF2b. Virology 287, 183–191 (2001).

10. Kuhn, J. H. et al. Reorganization and expansion of the nidoviral family arteriviridae. Archives of virology 161, 755–768 (2016).

11. Wissink, E. H. J. et al. Envelope protein requirements for the assembly of infectious virions of porcine reproductive and respiratory syndrome virus. J. Virol. 79, 12495–12506 (2005).

12. Dea, S., Gagnon, C. A., Mardassi, H., Pirzadeh, B. & Rogan, D. Current knowledge on the structural proteins of porcine reproductive and respiratory syndrome (PRRS) virus: comparison of the North American and European isolates. Arch. Virol. 145, 659–688 (2000).

13. Meng, X. J. Heterogeneity of porcine reproductive and respiratory syndrome virus: implications for current vaccine efficacy and future vaccine development. Vet. Microbiol. 74, 309–329 (2000).

14. Done, S. H., Paton, D. J. & White, M. E. Porcine reproductive and respiratory syndrome (PRRS): a review, with emphasis on pathological, virological and diagnostic aspects. Br. Vet. J. 152, 153–174 (1996).

15. Suárez, P. et al. Phylogenetic relationships of european strains of porcine reproductive and respiratory syndrome virus (PRRSV) inferred from DNA sequences of putative ORF-5 and ORF-7 genes. Virus Res. 42, 159–165 (1996).

16. Stadejek, T., Oleksiewicz, M. B., Potapchuk, D. & Podgorska, K. Porcine reproductive and respiratory syndrome virus strains of exceptional diversity in eastern Europe support the definition of new genetic subtypes. J. Gen. Virol. 87, 1835–1841 (2006).

17. Forsberg, R. et al. A Molecular Clock Dates the Common Ancestor of European-type Porcine Reproductive and Respiratory Syndrome Virus at More Than 10 Years before the Emergence of Disease. Virology 289, 174–179 (2001).

18. Forsberg, R. Divergence Time of Porcine Reproductive and Respiratory Syndrome Virus Subtypes. Mol. Biol. Evol. 22, 2131–2134 (2005).

19. Shi, M. et al. Phylogeny-based evolutionary, demographical, and geographical dissection of North American type 2 porcine reproductive and respiratory syndrome viruses. J. Virol. 84, 8700–8711 (2010).

20. Robinson, S. R., Abrahante, J. E., Johnson, C. R. & Murtaugh, M. P. Purifying selection in porcine reproductive and respiratory syndrome virus ORF5a protein influences variation in envelope glycoprotein 5 glycosylation. Infect. Genet. Evol.. 20, 362–368 (2013).

21. Rupasinghe, R. et al. Molecular Evolution of Porcine Reproductive and Respiratory Syndrome Virus Field Strains from Two Swine Production Systems in the Midwestern United States from 2001 to 2020. Microbiol. Spectr. 10, e0263421 (2022).

22. Zhao, Z.-Y. et al. Comparative analysis of newly identified rodent arteriviruses and porcine reproductive and respiratory syndrome virus to characterize their evolutionary relationships. Front. Vet. Sci. 10, 1174031 (2023).

23. Lu, Z. H. et al. Genomic variation in macrophage-cultured European porcine reproductive and respiratory syndrome virus Olot/91 revealed using ultra-deep next generation sequencing. Virol. J. 11, 42 (2014).

24. Hanada, K., Suzuki, Y., Nakane, T., Hirose, O. & Gojobori, T. The Origin and Evolution of Porcine Reproductive and Respiratory Syndrome Viruses. Mol. Biol. Evol. 22, 1024–1031 (2005).

25. Hilton, S. K. & Bloom, J. D. Modeling site-specific amino-acid preferences deepens phylogenetic estimates of viral sequence divergence. Virus Evol. 4, vey033 (2018).

26. Boni, M. F. et al. Evolutionary origins of the SARS-CoV-2 sarbecovirus lineage responsible for the COVID-19 pandemic. Nat. Microbiol. 5, 1408–1417 (2020).

27. Wu, Z. et al. Comparative analysis of rodent and small mammal viromes to better understand the wildlife origin of emerging infectious diseases. Microbiome 6, 178 (2018).

28. Plagemann, P. G. W. Porcine reproductive and respiratory syndrome virus: origin hypothesis. Emerg. Infect. Dis. 9, 903–908 (2003).

29. McMullen, L. K. Berkshire Swine Production and Marketing. (2006).

30. Smith, J. L. Animal Farms. in Works in Progress: Plans and Realities on Soviet Farms, 1930-1963 (Yale University Press, 2014). doi:10.12987/yale/9780300200690.003.0004

31. Zwart, M. P. & Elena, S. F. Matters of Size: Genetic Bottlenecks in Virus Infection and Their Potential Impact on Evolution. Annu. Rev. Virol. 2, 161–179 (2015).

32. McCrone, J. T. & Lauring, A. S. Genetic bottlenecks in intraspecies virus transmission. Curr. Opin. Virol. 28, 20–25 (2018).

33. Tamura, K. et al. MEGA5: molecular evolutionary genetics analysis using maximum likelihood, evolutionary distance, and maximum parsimony methods. Mol. Biol. Evol. 28, 2731–2739 (2011).

34. Kosakovsky Pond, S. L., Posada, D., Gravenor, M. B., Woelk, C. H. & Frost, S. D. W. GARD: a genetic algorithm for recombination detection. Bioinformatics 22, 3096–3098 (2006).

35. Lam, H. M., Ratmann, O. & Boni, M. F. Improved Algorithmic Complexity for the 3SEQ Recombination Detection Algorithm. Mol. Biol. Evol. 35, 247–251 (2018).

36. Martin, D. P., Murrell, B., Golden, M., Khoosal, A. & Muhire, B. RDP4: Detection and analysis of recombination patterns in virus genomes. Virus Evol. 1, vev003 (2015).

37. Pond, S. L. K., Frost, S. D. W. & Muse, S. V. HyPhy: hypothesis testing using phylogenies. Bioinformatics 21, 676–679 (2005).

38. Murrell, B. et al. FUBAR: A Fast, Unconstrained Bayesian AppRoximation for Inferring Selection. Mol. Biol. Evol. 30, 1196–1205 (2013).

39. Smith, M. D. et al. Less Is More: An Adaptive Branch-Site Random Effects Model for Efficient Detection of Episodic Diversifying Selection. Mol. Biol. Evol. 32, 1342–1353 (2015).

40. Drummond, A. J., Suchard, M. A., Xie, D. & Rambaut, A. Bayesian phylogenetics with BEAUti and the BEAST 1.7. Mol. Biol. Evol. 29, 1969–1973 (2012).

41. Raftery, A. E., Niu, X., Hoff, P. D. & Yeung, K. Y. Fast Inference for the Latent Space Network Model Using a Case-Control Approximate Likelihood. J. Comput. Graph. Stat. 21, 901–919 (2012).

42. Ward, B. J. & van Oosterhout, C. hybridcheck: software for the rapid detection, visualization and dating of recombinant regions in genome sequence data. Mol. Ecol. Resour. 16, 534–539 (2016).

43. Malinsky, M., Matschiner, M. & Svardal, H. Dsuite - Fast D-statistics and related admixture evidence from VCF files. Mol. Ecol. Resour. 21, 584–595 (2021).

44. Hubisz, M. & Siepel, A. Inference of Ancestral Recombination Graphs Using ARGweaver. in Statistical Population Genomics (ed. Dutheil, J. Y.) 231–266 (Springer US, 2020). doi:10.1007/978-1-0716-0199-0_10

45. Griffiths, R. C. & Marjoram, P. An ancestral recombination graphitle. in Progress in population genetics and human evolution 257–270 (Springer, 1997).

46. Morel, B., Kozlov, A. M., Stamatakis, A. & Szöllősi, G. J. GeneRax: A Tool for Species-Tree-Aware Maximum Likelihood-Based Gene Family Tree Inference under Gene Duplication, Transfer, and Loss. Mol. Biol. Evol. 37, 2763–2774 (2020).

47. Penel, S., Menet, H., Tricou, T., Daubin, V. & Tannier, E. Thirdkind: displaying phylogenetic encounters beyond 2-level reconciliation. Bioinformatics 38, 2350–2352 (2022).

