## Supplementary material for "Rodent Origins and Human-Mediated Evolutionary Dynamics of PRRSV": Supplementary Figures.docx

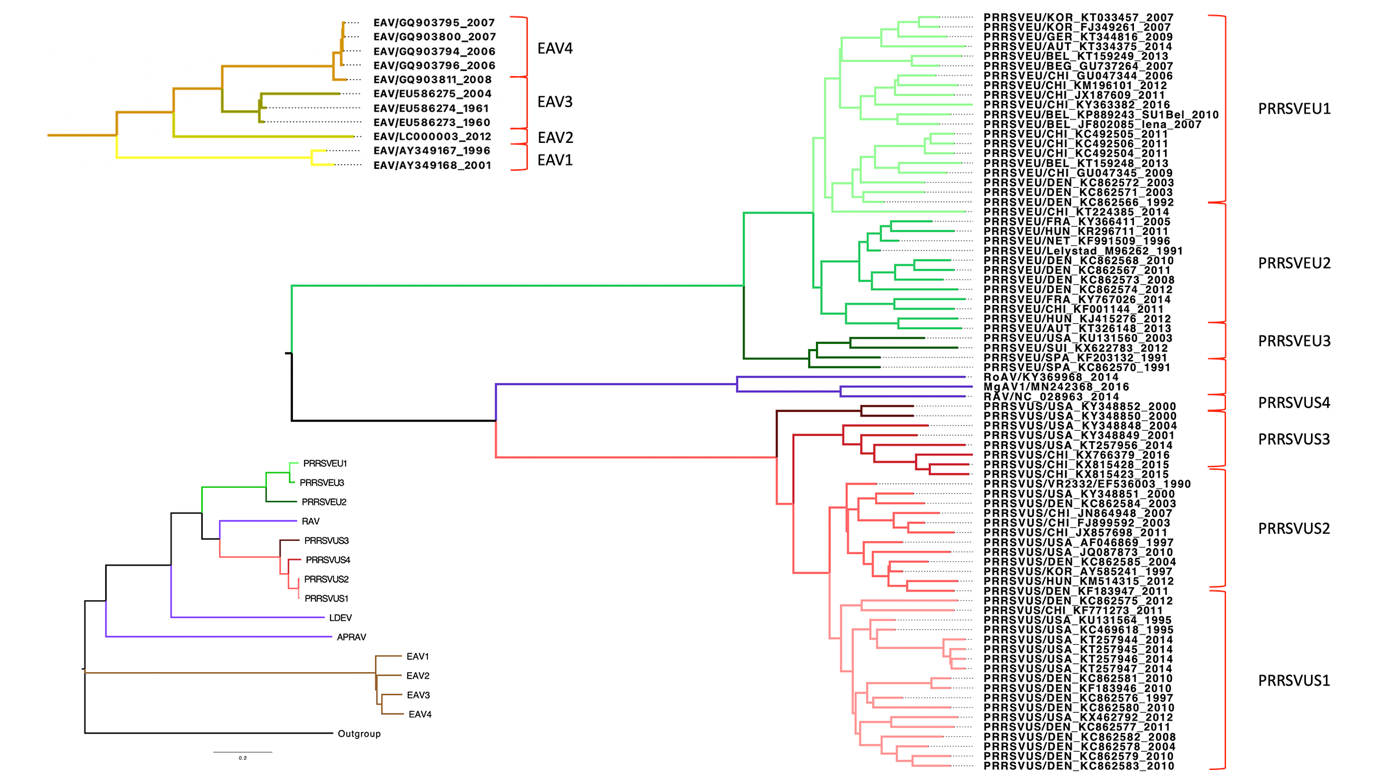

**Supplementary Figure 1. MCA Tree.**

Viral species are represented in different colors, while clades within each species are shaded using varying gradients of the primary color assigned to the species. These color gradients are consistent with those used in subsequent analyses and are directly compatible with this tree.

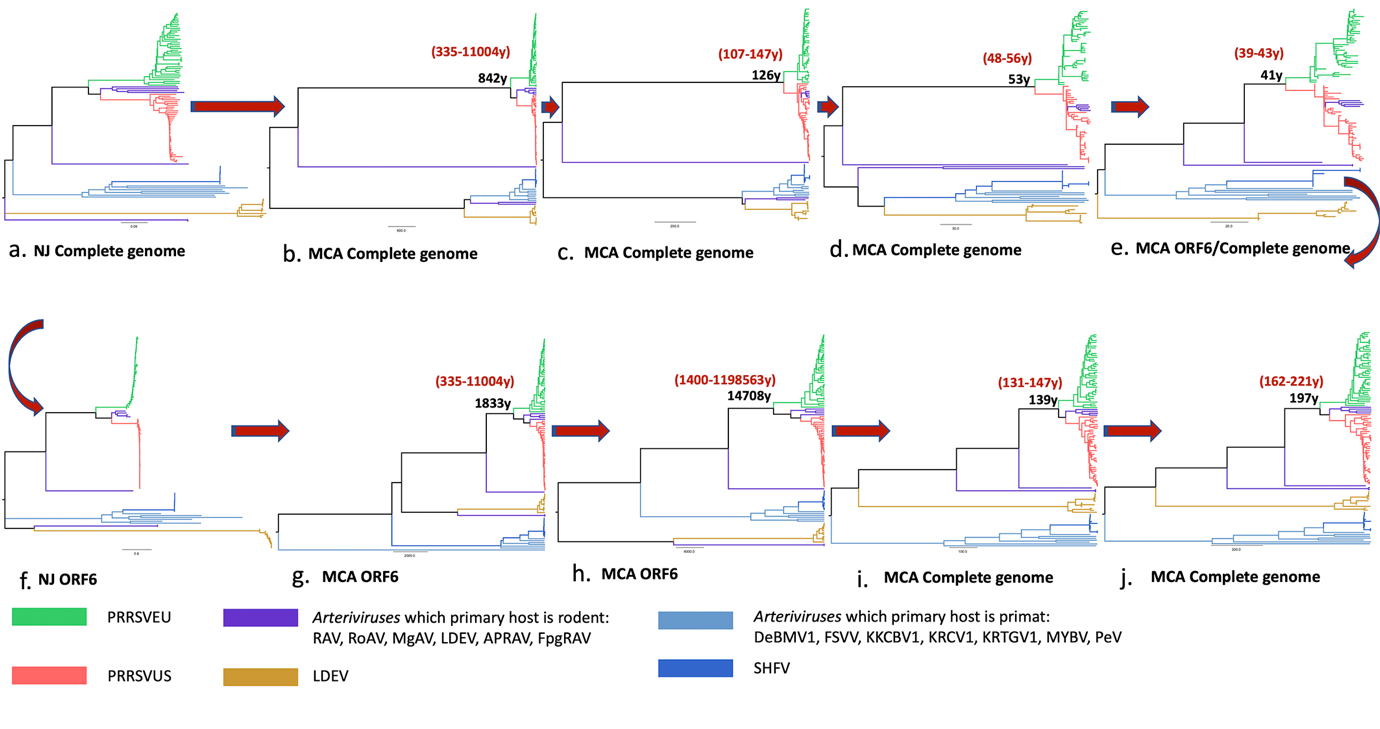

**Supplementary Figure 2. Phylogenetic Trees.**

The figure presents the results of the phylogenetic analysis: (a) An Neighbor-Joining (NJ) tree was constructed using the complete genome sequences of Arteriviruses with the Tamura-Nei, substitution model. (b) An MCA tree was constructed using the complete genome sequences of Arteriviruses with the GTR + Г4 + I, substitution model, an uncorrelated relaxed exponential molecular clock, and a coalescent growth tree prior. (c) Another MCA tree was built using the same dataset but with the YANG96 + Г4 + I substitution model, an uncorrelated relaxed exponential molecular clock, and a coalescent exponential growth tree prior. (d) MCA trees were generated for partitions representing individual ORFs from the complete genome dataset. Each partition used the YANG96 + Г4 + I substitution model and an uncorrelated relaxed exponential molecular clock. The coalescent exponential growth tree prior was set for the complete genome partition, with other ORF partitions linked to it; (e) Similarly, MCA trees were created with ORF partitions linked to ORF6, using the same substitution model and clock settings. (f) A Neighbor-Joining (NJ) tree was computed for the ORF6 partition dataset, excluding two sequences with gaps due to incomplete sequencing. The gaps were minor and did not affect the overall analysis. (g) An MCA tree was constructed using the ORF6 partition with the YANG96 + Г4 + I substitution model, an uncorrelated relaxed exponential molecular clock, and a coalescent growth tree prior. (h) An MCA tree was constructed using the ORF6 partition with the YANG96 + Г4 + I substitution model, an uncorrelated relaxed exponential molecular clock, and a Byesian skyline coalescent tree prior. (i) An MCA tree was developed using codons from the complete genome dataset, excluding regions under positive or negative selection (based on FEL analysis) or identified as recombinant (via 3SEQ analysis). YANG96 + Г4 + I substitution model, an uncorrelated relaxed exponential molecular clock, and a coalescent growth tree prior were applied. Tree priors for PRRSV-1 and PRRSV-2 were set according to analyses focused on PRRSV and Rodent/Rat arterivirus clades. (j) A similar analysis to (h) was conducted using a Bayesian skyline coalescent tree prior instead of the growth prior. Tree priors for PRRSV-1 and PRRSV-1 were similarly refined. . The colors used to represent species are consistent with those in Suppl.fig 1. For each MCA tree, the TreeHeight value is shown in black, and the 95% HPD (highest posterior density) interval is indicated in red.

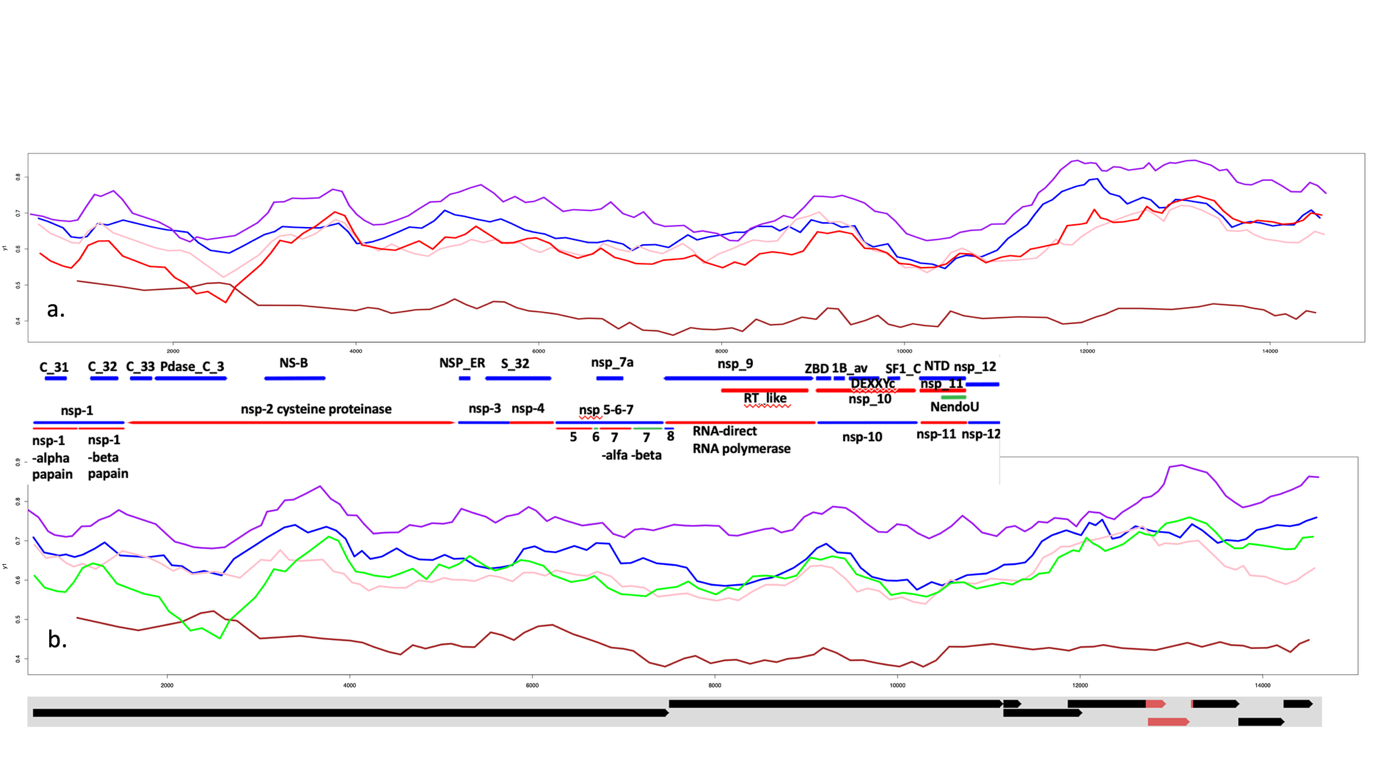
 **Supplementary Figure 3. Results of DInvestigate Analysis.**

**a.** Introgression of PRRSV-2, RAV, LDEV, APRAV, and EAV into PRRSV-1, subtype 1. **b.** Introgression of PRRSV-1, RAV, LDEV, APRAV, and EAV into PRRSV-2. Legend: Brown line = EAV, Blue line = LDEV, Purple line = RAV, Pink line = APRAV, Green line = PRRSV-2, Red line = PRRSV-1.

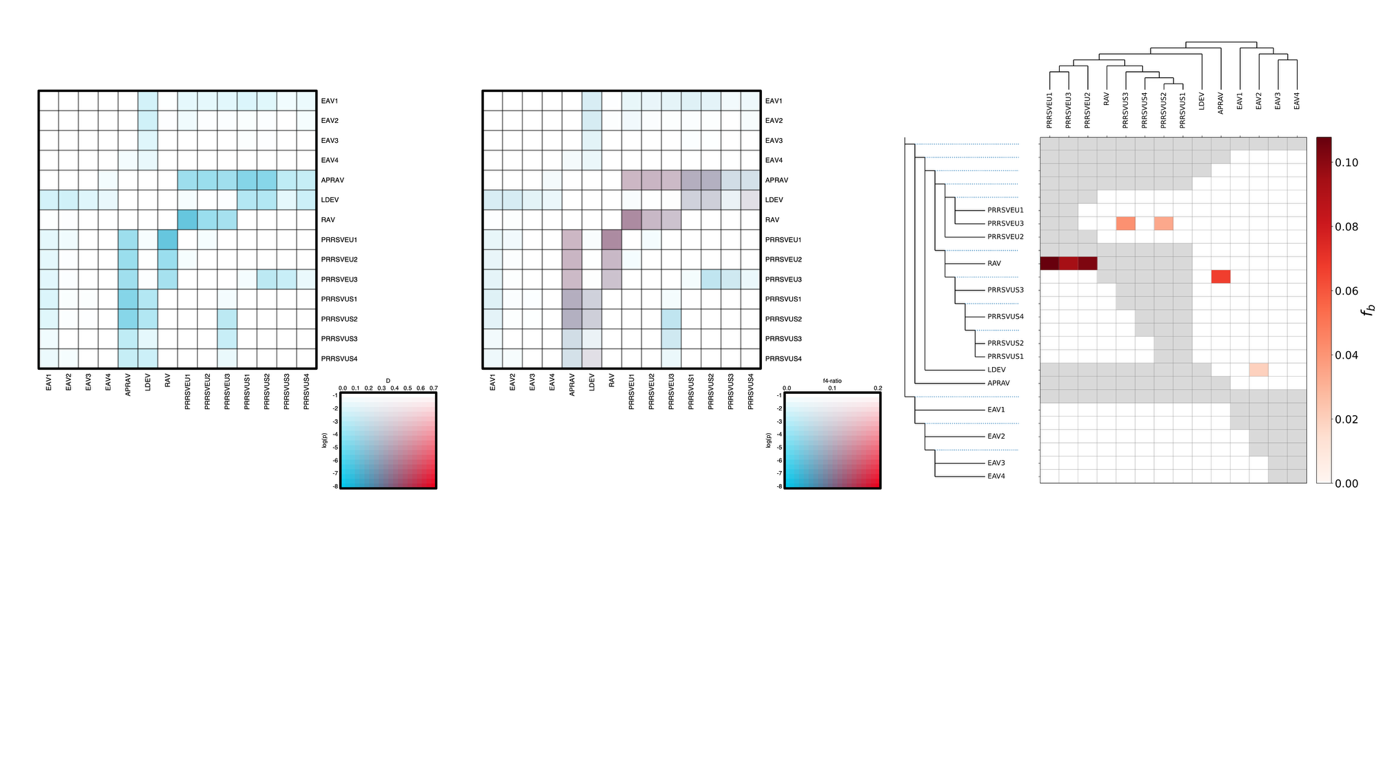

**Supplementary Figure 4: Results from Dsuite Analysis. a: D-matrix (D-statistics).**

This heatmap represents pairwise D-statistics between taxa. D-statistics measure deviations from expected allele frequencies under a strict tree-like model of evolution, identifying potential introgression. Blue-shaded cells indicate significant deviations, with stronger blue tones reflecting higher statistical significance. b: f4-ratio Matrix. This heatmap visualizes the f4-ratio, which quantifies the proportion of introgressed material between pairs of taxa. Stronger purple shading indicates a higher proportion of introgressed genetic material. c: fD Matrix with Phylogenetic Tree. This matrix incorporates fD values, a refined statistic for detecting introgression, plotted alongside a phylogenetic tree. Higher fD values (red shading) indicate stronger introgression signals. Taxa relationships are represented on the left as a phylogenetic tree.

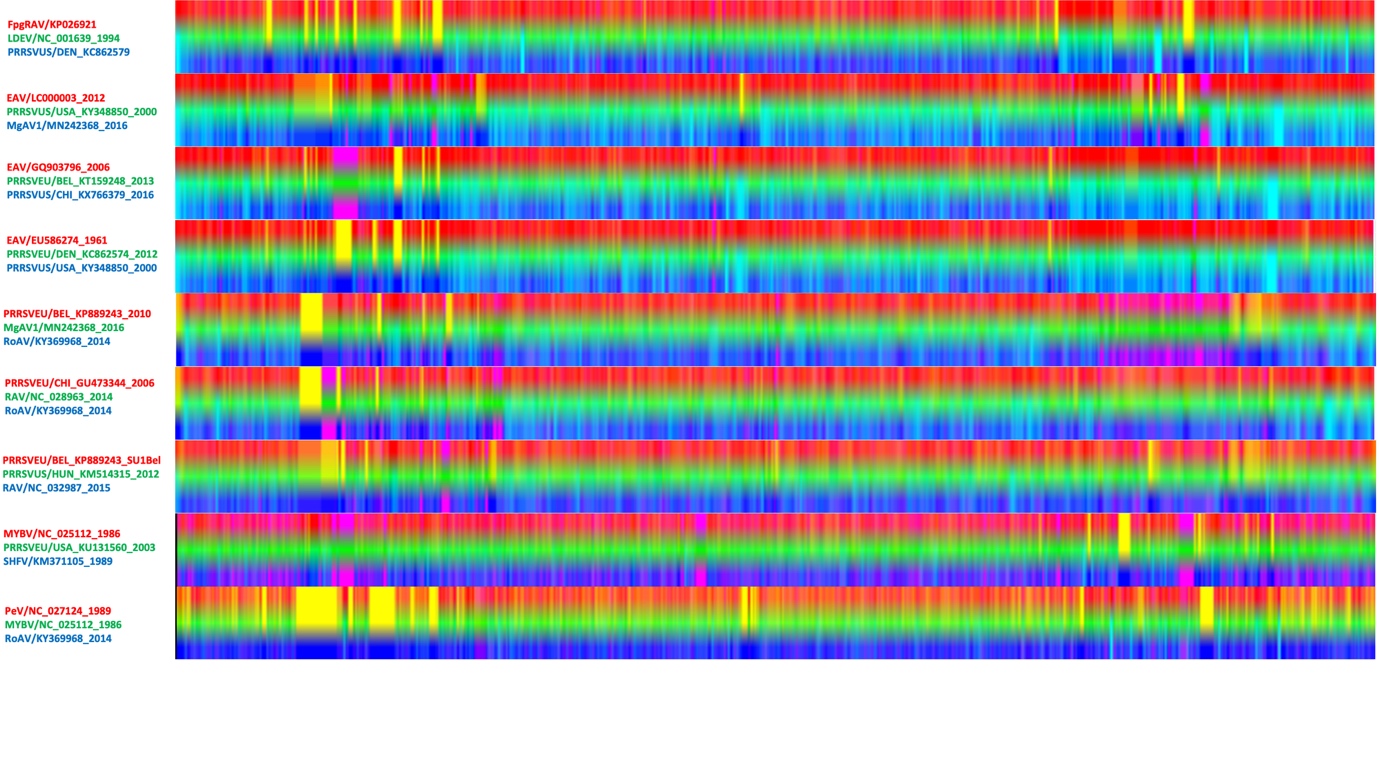

**Supplementary Figure 5. HybridCheck Analysis**

The x-axis represents nucleotide positions, while the y-axis corresponds to the taxa triplets of sequences tested. Red, green, and blue colors represent genetic similarity or the background phylogenetic relationships, whereas yellow, cyan, and pink highlight regions of introgression.

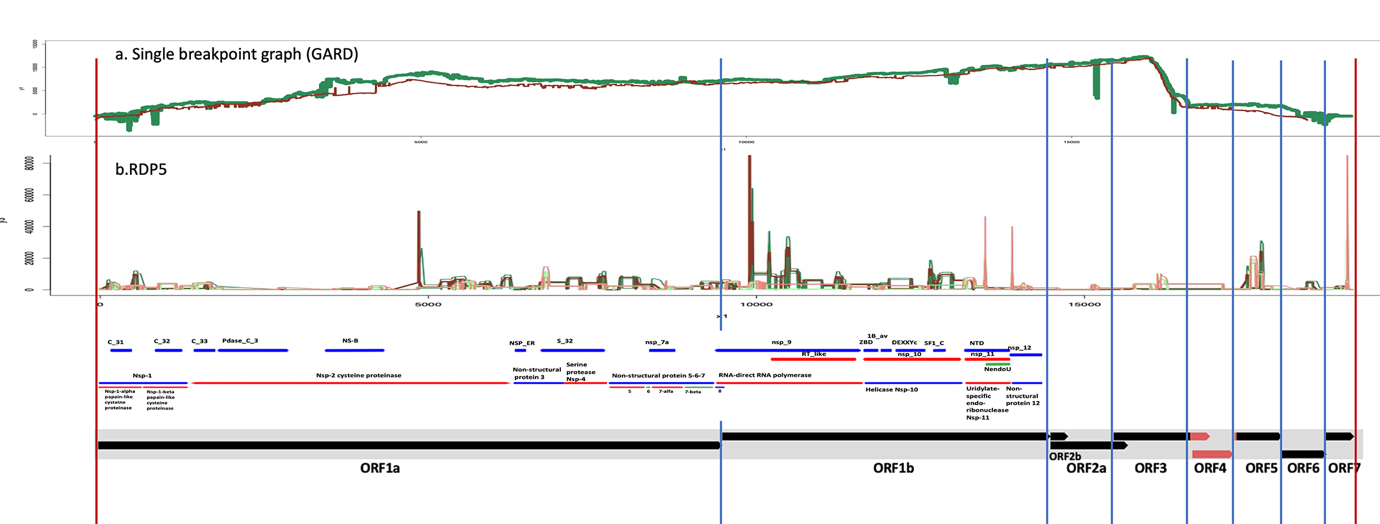

**Supplementary Figure 6. Visual Representation of Results fo Recombination test.**

This figure provides a comprehensive summary of the results from various analyses: a: SBP Graph. AIC (Akaike Information Criterion) values are plotted for the AV_cod dataset (green) and the PRRSV_cod dataset (red). b: RDP5. Confidence intervals (CIs) are shown for the AV_cod (green) and PRRSV_cod (red) datasets. Light green/red represents the 95% CI, while dark green/red represents values beyond the 95% CI.

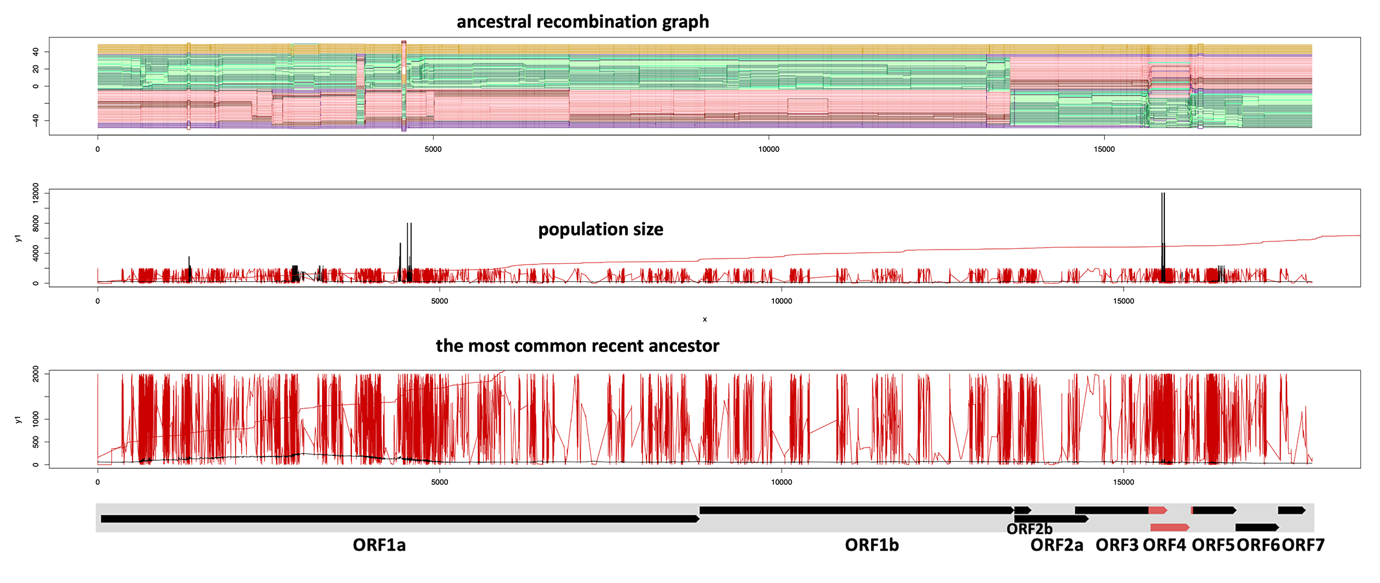

**Supplementary figure 7. Ancestral Recombination Graph.**

a. Ancestral Recombination Graph: This panel displays an ancestral recombination graph (ARG), illustrating how recombination events shape the evolutionary history of the sequences. The y-axis represents potential recombination lineages, while the x-axis indicates nucleotide positions along the genome. b. Population Size: The population size is inferred across the genome, with peaks indicating regions of higher effective population size. The x-axis represents genomic positions, and the y-axis shows population size estimates. c. Most Recent Common Ancestor: This panel depicts the most recent common ancestor inferred for each genomic region. Peaks correspond to higher ancestral coalescence times. The x-axis represents nucleotide positions, and the y-axis indicates coalescent times.

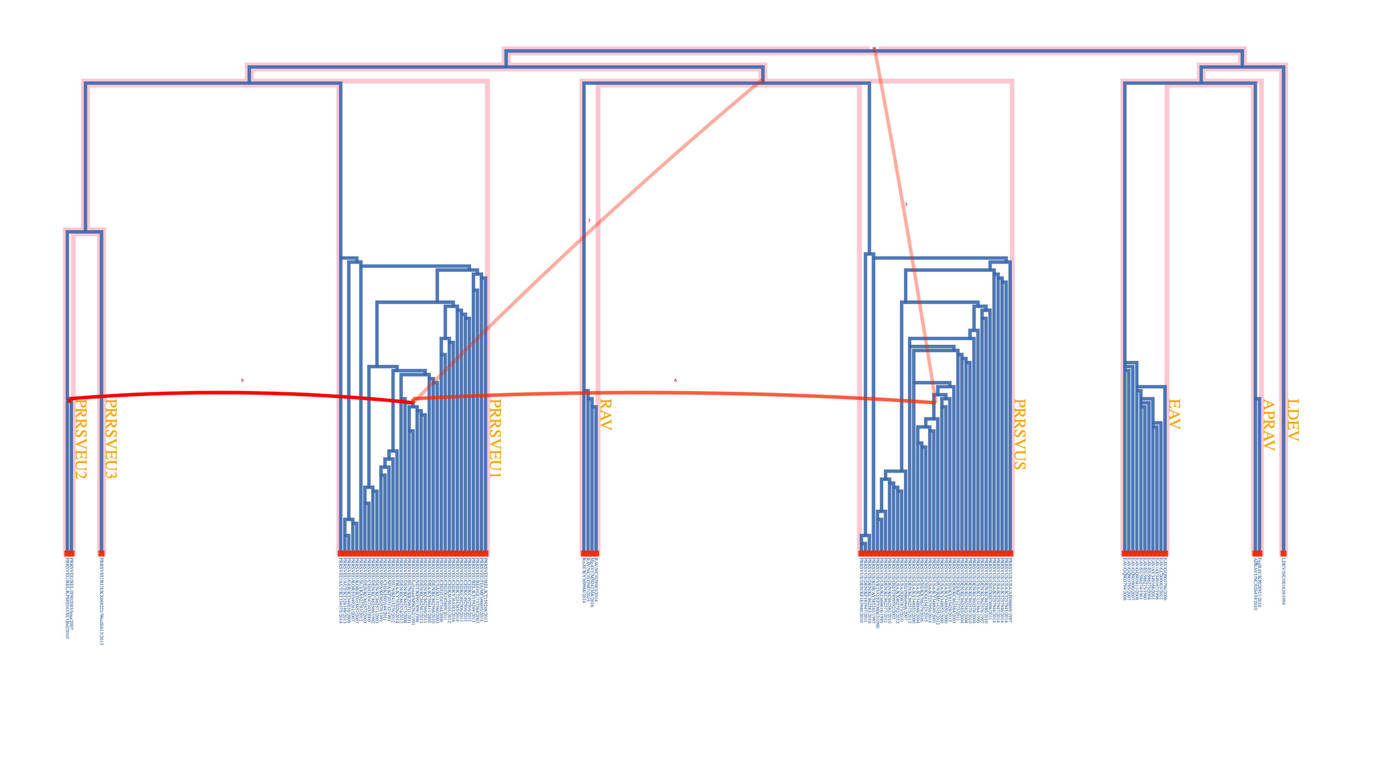

**Supplementary Figure 8. ARG tree. Analysis of GeneRax Output visualised in ThirdKind for ORFs.**

Phylogenetic Trees (Blue Lines): The blue trees represent the inferred phylogenetic structure for each clade, including PRRSV-1 subtype 1, PRRSV-1 subtype 2, PRRSV-1 subtype 3, PRRSV-2, RAV, EAV, APRAV, and LDEV. These trees depict hierarchical evolutionary relationships within and between groups of arteriviruses. Horizontal Red Arcs: The red arcs represent horizontal gene transfer (HGT) events between specific clades or taxa. Pink connections reinforce the evidence of HGT between specific nodes or branches, indicating potential shared ancestry or gene flow between taxa.

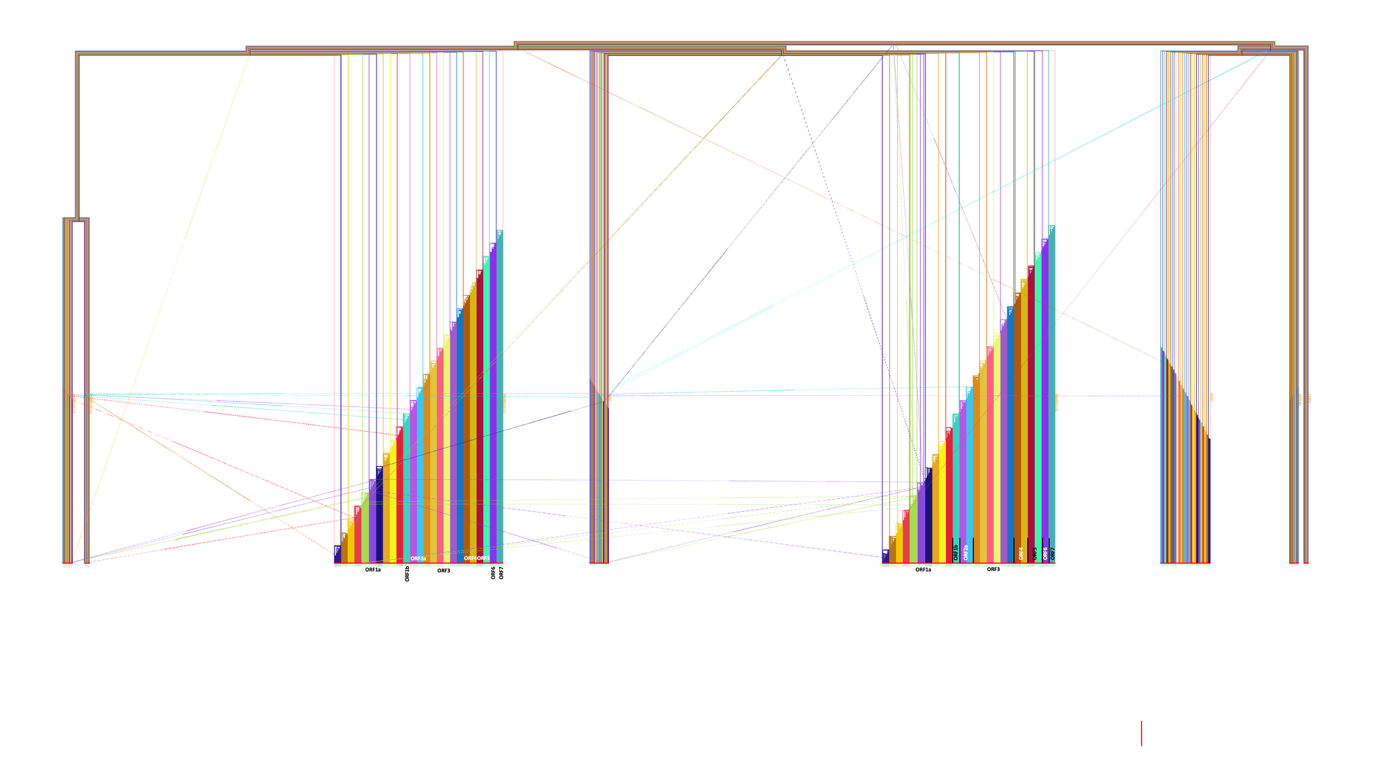
 **Supplementary Figure 9. ARG tree. Analysis of GeneRax Output visualised in ThirdKind for recombinant blocks.**

Phylogenetic Trees (Blue Lines): The different coloured trees represent the inferred phylogenetic structure for, including PRRSV-1 subtype 1, PRRSV-1 subtype 2, PRRSV-1 subtype 3, PRRSV-2, RAV, EAV, APRAV, and LDEV. These trees depict hierarchical evolutionary relationships within and between groups of arteriviruses. Horizontal different coloured Arcs (HGT Events): The arcs represent horizontal part of gene transfer (HGT) events between specific clades or taxa. Connections reinforce the evidence of HGT between specific nodes or branches, indicating potential shared ancestry or gene flow between taxa.

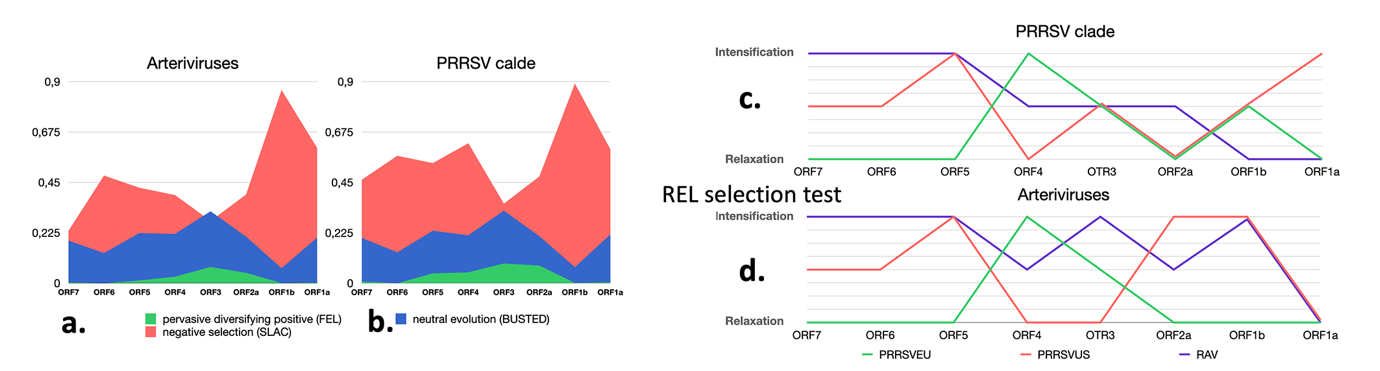

**Supplementary Figure 10. Selection pressuer on ORFs**

a. Selection pressure on encoding regions of the Arterivirus clade. B. Selection pressure on encoding regions of the PRRSV clade. c. Selection trends in PRRSV. d. Selection trends across the complete Arterivirus dataset.

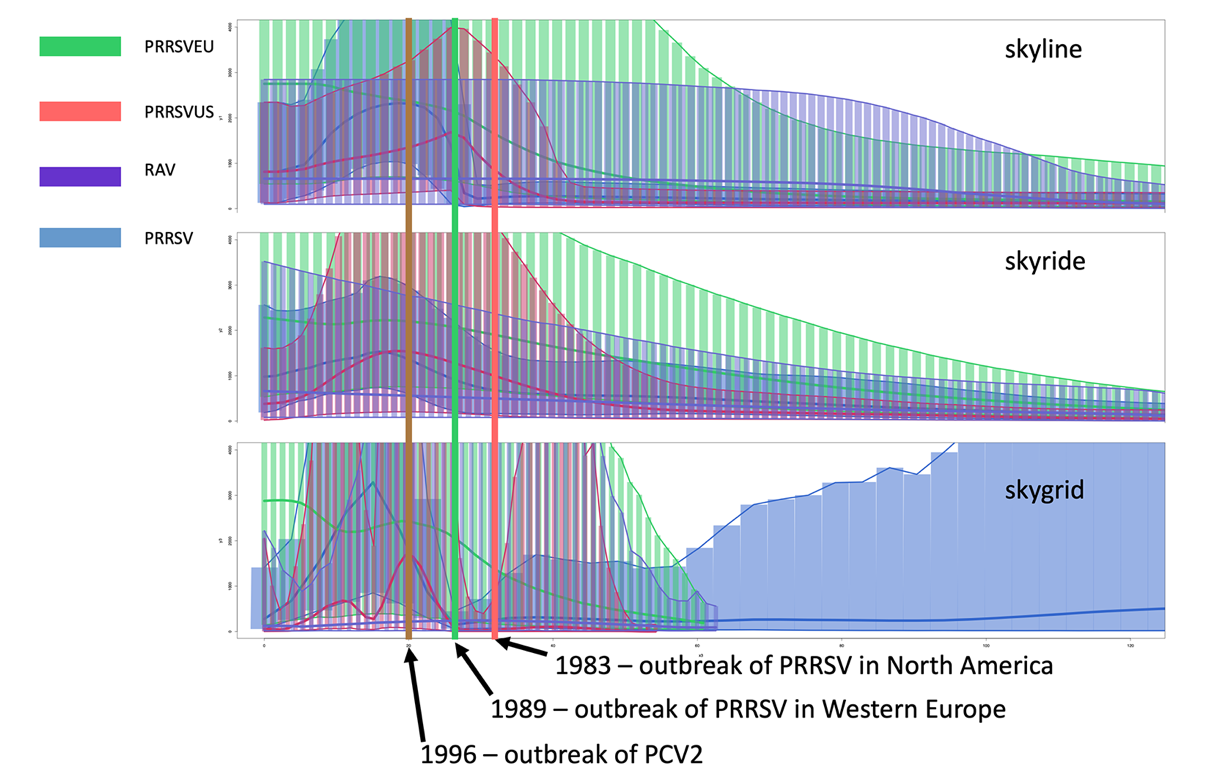

**Supplementary Figure 11. Skyplots.**

a. Skyline, b. Skyride, and c. Skygrid Analyses applied to different datasets. Colours and Representations: Blue: All sequences (PRRSV + Rodent arteriviruses combined). Red: PRRSV-2, Green: PRRSV-1, Violet: RAV.

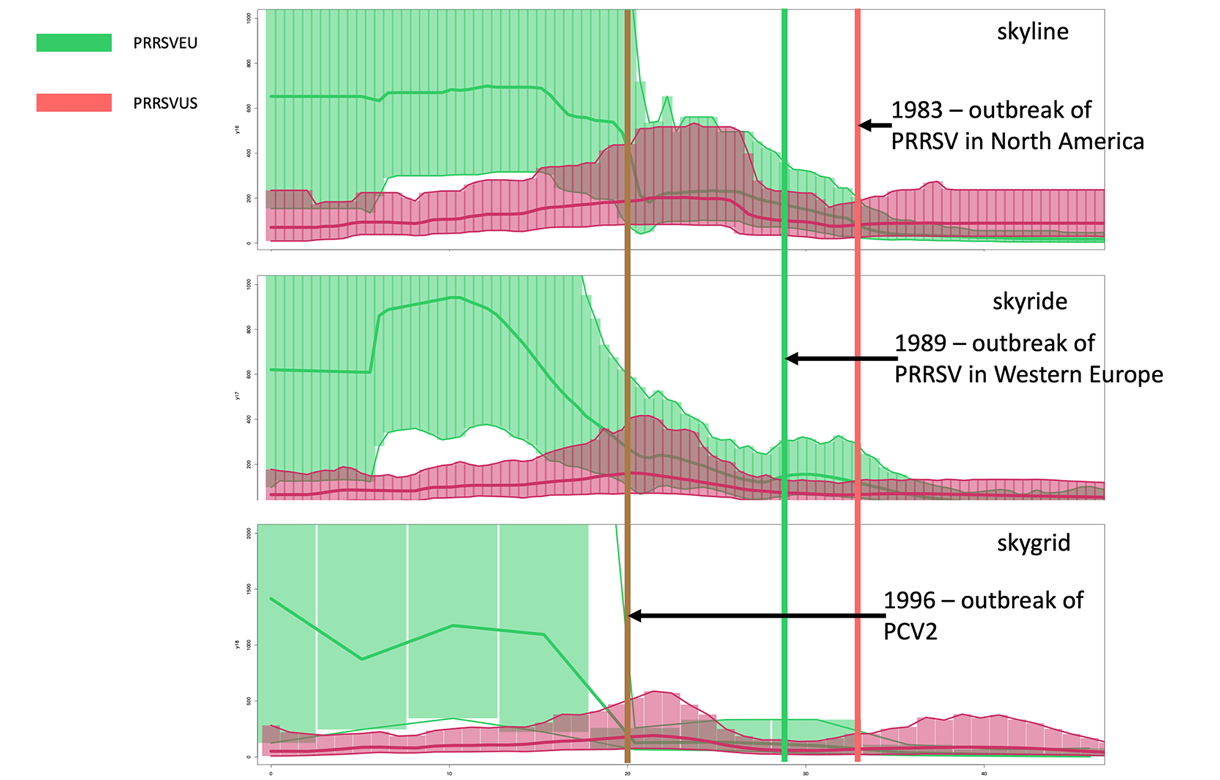

**Supplementary Figure 12. Skyplots of PRRRSV**

Adjusted Skyplot Analyses a. Skyline, b. Skyride, and c. Skygrid. This analysis removes codons under positive selection and recombinant regions, focusing on the neutral evolutionary signal. Colors: PRRSV-1, Red: PRRSV-2.

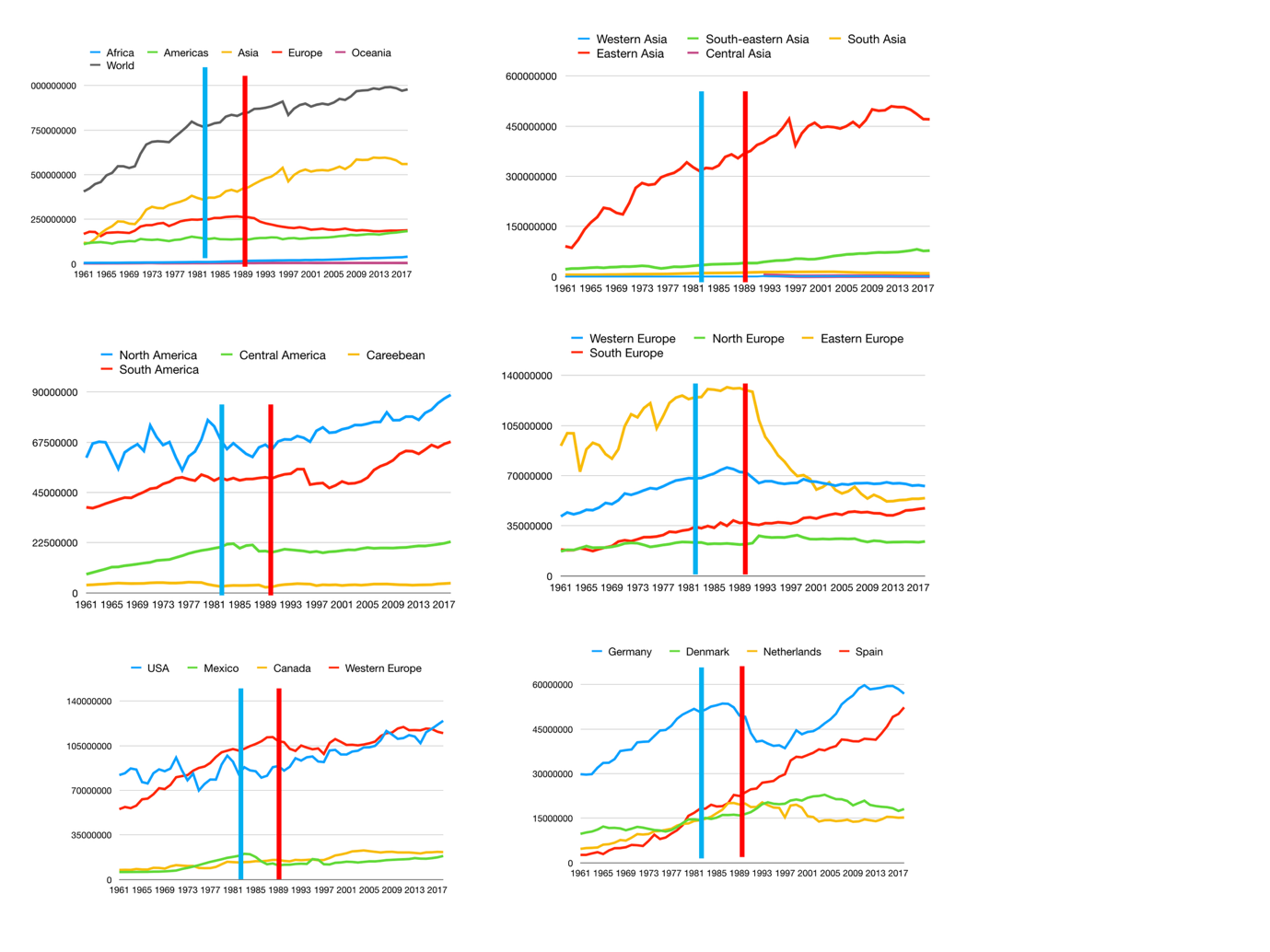

**Supplementary Figure 13. Pig production worldwide in correlation with PRRSV outbreaks.**

The blue horizontal bar indicates the PRRSV outbreak in Southern U.S. states in 1983–1984, The red horizontal bar highlights the PRRSV outbreak in Lower Saxony, Germany, in 1989. a. Continents – Pig production density by continent. b. Regions in Asia – Distribution and trends in pig production across key regions in Asia. c. Regions in the Americas – Pig production trends across North, Central, and South America. d. Regions in Europe – Pig production trends in various European regions. e. North American states vs. Western Europe – Comparative trends in pig production between major North American states and Western European countries. f. Top-producing states in Europe – Pig production density in the largest pig-producing states in Europe. Data Source: FAO (Food and Agriculture Organization).

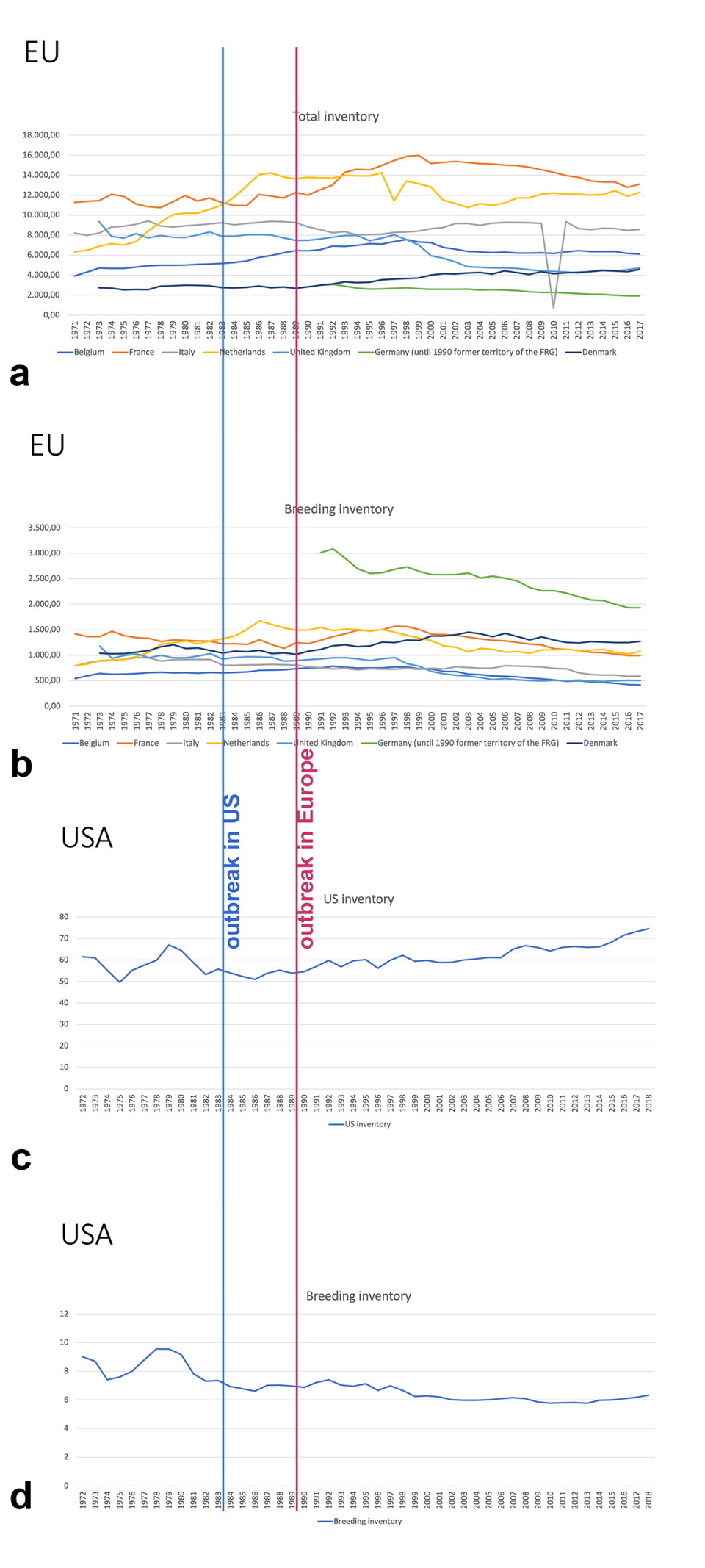

**Supplementary Figure 14. Pig inventory.**

This image presents trends in pig inventory, focusing on total inventory and breeding inventory for Europe (EU) and the United States (USA) from the 1970s to 2019. by EUROSTATA and USDA for a. EU Total Inventory, b. EU Breeding Inventory; c, US Total Inventory; d. US Breeding Inventory.

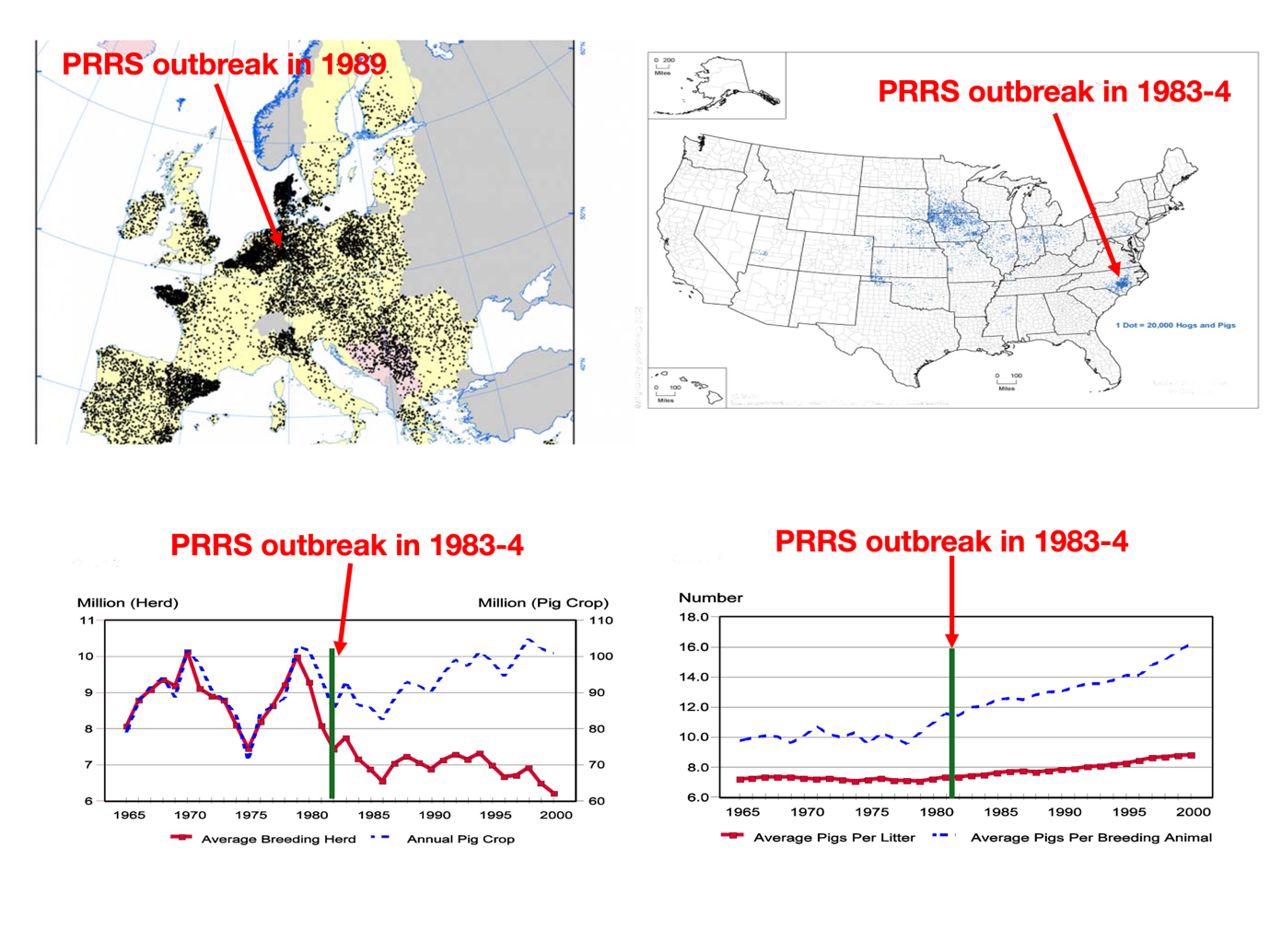

**Supplementary Figure 14. Pig demography**

a. Pig population density in Europe – each dot represents 1,000 pigs. b. Pig population density in the United States – each dot represents 20,000 pigs. c. Pig production trends in the United States – correlation between the average breeding herd size (red solid line) and the annual pig crop (blue dashed line) from 1965 to 2000. The PRRS outbreak in 1983–84 is highlighted. d. Reproductive performance trends in the United States – shows changes in the average pigs per litter (red solid line) and average pigs per breeding animal (blue dashed line) from 1965 to 2000. The PRRS outbreak in 1983–84 is marked for context. Data Source: Figure 17a from Eurostat, 17b-d from USDA.
