## Supplementary material for "Rodent Origins and Human-Mediated Evolutionary Dynamics of PRRSV": Supplementary Table Legends.docx

**Table S1 Model testing for PRRSV phylogenetic dating analyses.**

Comparison of substitution models, molecular‐clock models and tree priors fitted to whole-genome PRRSV datasets (entries PRRSV1–PRRSV22) and to ORF6, with corresponding model IDs. For each run the table lists: codon (SRD06 or Yang96) or nucleotide (GTR) model; treatment of base frequencies (empirical vs estimate); clock model (strict; relaxed log = lognormal; relaxed exp = exponential; relaxed gamma); and coalescent prior (Const = constant population size; Exponential growth; Skyline). Model fit is summarized by AICM (Akaike Information Criterion through Markov chains; lower is better) and its standard error; posterior likelihood summary statistics (mean, standard deviation, standard error, variance); and ESS (effective sample size) as a convergence diagnostic (larger values indicate better mixing). Blank cells denote values not reported or not applicable.

**Table S2 Recombination screen using 3SEQ.**

Results of the 3SEQ scan for recombination across arteriviruses and PRRSV. The workbook contains four sheets: **AV_rec**(arterivirus-wide screen by genomic region), **PRRSV_rec** (PRRSV-only screen), and summary lists of sequences with **long recombinant tracts** (**AV_long_rec**, **PRRSV_long_rec**). For each putative event, the table reports the **candidate recombinant** (**C_ACCNUM**), the two **putative parents** (**P_ACCNUM** columns), 3SEQ test statistics (**m, n, k**), the exact **P value** (p) and the maximum P across breakpoint choices (**[p_max]**), an indicator of **homoplasy support** (**HS?**), **log(p)**, and **down-sampling summaries** (**DS(p)**; mean/variance as shown). The **minimum recombinant-fragment length** and inferred **breakpoint coordinates** (primary and alternative intervals in subsequent columns) are given in **alignment positions**. Accession labels follow the dataset convention (lineage/iso-code_accession_year). Blank entries indicate partitions with no detectable signal. Significance thresholds and post-processing filters (masking, multiple testing, parental relatedness and duplicate removal) are described in **Methods**.

**Table S3 Selection analyses across PRRSV and related arteriviruses.**

Summary of codon‐based selection tests applied to whole genomes and individual ORFs. The workbook aggregates gene-level, site-level and branch/branch-site results; sheet names follow the corresponding method (e.g., FEL, MEME, FUBAR, BUSTED, aBSREL, and per-gene summaries). For each analysis we report dataset/gene, alignment length, number of sequences, model specification, test statistic (e.g., LRT or Bayes factor), ω (dN/dS) estimates, p or q values (FDR where applicable), and counts/lists of sites under pervasive (FEL/FUBAR) or episodic (MEME) diversifying selection. Branch/branch-site results (aBSREL/BUSTED) include the tested lineage(s), fitted rate classes, proportion under selection and support values. Site coordinates are given in codon positions and, where provided, mapped to reference genomes (e.g., Lelystad/VR-2332). Analyses were run on recombination-aware alignments (masked or partitioned as described in Methods), and significance was defined by method defaults (typically p < 0.05 or q < 0.1 for site-wise tests) unless stated otherwise. Blank cells indicate metrics not applicable to a given method.

**Table S4 Historical changes in pig production systems relevant to PRRSV emergence.**

Chronological summary of major changes in pig production and breeding practices that altered herd structure, contact patterns and gene flow among farms. Entries list the year and a brief description of the innovation or management shift, including: the move from outdoor to confined, stage-structured production (breeding, gestation, farrowing, nursery, grower, finisher); the development of pyramidal breeding systems with nucleus herds for genetic improvement; the introduction and subsequent widespread adoption of artificial insemination (AI) and cryopreserved boar semen; the emergence of medicated early weaning (MEW), age-segregated rearing (ASR) and related multi-site weaning systems; the large-scale roll-out of + multi-site production in North America; and the establishment of large, mechanized collective pig farms in the Soviet Union. Together, these milestones trace the intensification and restructuring of pig production that plausibly shaped the ecological context for PRRSV amplification and spread.

**Table S5 ORF partitions and genome coordinate ranges by species.**

Partition scheme used for downstream analyses, listing (for each species) the **number of sequences** and the **genome nucleotide ranges** defining **ORF1a, ORF1b, ORF2ab, ORF3, ORF4, ORF5, ORF6** and **ORF7.** Coordinate intervals are **1-based, inclusive** and correspond to the species-specific reference annotations employed in this study; **“ORF2ab”** denotes merged ORF2a/ORF2b where applicable. These partitions were used for alignment masking/partitioning, recombination screening, phylogenetic inference, and selection analyses as described in **Methods.** Species abbreviations are provided below the table (e.g., **APRAV, EAV, LDEV, PRRSV-1/2, RAV, SHFV, etc**.).
