## Supplementary material for "Rodent Origins and Human-Mediated Evolutionary Dynamics of PRRSV": Table_S1_Model_test.docx

**Table S1 Model testing for PRRSV phylogenetic dating analyses.**

|  | Model | | | | AICM | | Likelihood | | | | | |
| --- | --- | --- | --- | --- | --- | --- | --- | --- | --- | --- | --- | --- |
| ID | Codon model | Base frequencies | Molecular clock | Prior | AICM | Stderr | | Mean | Stdev | Sterr | Variance | ESS |
| PRRSV1 | SRD06 | estimate | strict | Const | 743420,284 | -0,127 | | -371633,27 | 0,0824 | 8,7679 | 76,876 | 11335 |
| PRRSV2 | SRD06 | empirical | strict | Const | 743458,405 | -0,327 | | -371653,5 | 0,0812 | 8,7009 | 75,7055 | 11491 |
| PRRSV3 | Yang96 | estimate | strict | Const | 741613,664 | -0,948 | | -370654,49 | 0,2212 | 12,3426 | 152,3386 | 3112,15 |
| PRRSV4 | **Yang96** | **empirical** | strict | Const | 741658,961 | -0,409 | | -370675,33 | 0,3264 | 12,3999 | 153,7587 | 1443,1075 |
| PRRSV5 | Yang96 | estimate | log | Const | 738040,551 | -1,171 | | -368771,91 | 0,5126 | 15,7597 | 248,3691 | 945,3449 |
| PRRSV6 | Yang96 | estimate | **exp** | Const | 737976,258 | -0,885 | | -368759,68 | 0,4954 | 15,1145 | 228 | 930,9582 |
| PRRSV7 | Yang96 | estimate | gamma | Const | 738002,668 | -1,397 | | -368762,77 | 0,3984 | 15,4455 | 238,5637 | 1502,6301 |
| PRRSV9 | Yang96 | estimate | fixed | Const | 741589,841 | -0,453 | | -370638,86 | 0,2891 | 12,4926 | 156,0643 | 1867,4259 |
| PRRSV10 | Yang96 | estimate | strict/log | Const | 738672,420 | -1,489 | | -368653,34 | 0,8056 | 26,1319 | 682,8744 | 1052,1809 |
| PRRSV11 | Yang96 | estimate | log/log | Const | 733607,768 | -2,049 | | -366164,43 | 0,9716 | 25,2874 | 639,4539 | 677,3436 |
| PRRSV12 | Yang96 | estimate | EXP/strict | Const | 737976,258 | -0,885 | | -368759,68 | 0,4954 | 15,1145 | 228 | 930,9582 |
| PRRSV13 | Yang96 | estimate | EXP/log | Const | 734626,356 | -1,668 | | -367003,99 | 0,7009 | 17,5895 | 309 | 629,8283 |
| PRRSV14 | Yang96 | estimate | **EXP/exp** | Const | 734608,929 | -1,177 | |  |  |  |  |  |
| PRRSV15 | **Yang96** | **estimate** | **EXP/gamma** | **Const** | **734604,075** | **-1639** | | **-366983,03** | **0,9516** | **17,862** | **319,0514** | **352,3552** |
| PRRSV16 | **Yang96** | **estimate** | **exp/strict** | **Const** | **737976,258** | **-0,885** | | **-368759,68** | **0,4954** | **15,1145** | **228** | **930,9582** |
| PRRSV17 | **Yang96** | **estimate** | **exp/log** | **Const** | **733616,484** | **-1,593** | | **-366171,45** | **0,719** | **25,2347** | **636,7898** | **1231,6967** |
| PRRSV18 | **Yang96** | **estimate** | **exp/exp** | **Const** | **733322,752** | **-1,94** | | **-366106,91** | **0,5353** | **23,5472** | **554,4685** | **1935,2228** |
| PRRSV19 | **Yang96** | **estimate** | **exp/exp** | **Exponential** | **733262,370** | **-2,129** | | **-366080,99** | **0,4751** | **23,4561** | **550,1908** | **2437,4179** |
| PRRSV22 | **Yang96** | **estimate** | **exp/exp** | **Skyline** | **733278,260** | **-3,679** | | **-366082,43** | **0,564** | **23,5946** | **556,7042** | **1750,118** |
| ORF6 | **Yang96** | **estimate** | **exp** | **Exponential** | **12937,908** | **-0,197** | |  |  |  |  |  |
| ORF6 | GTR | estimate | exp | Exponential | 13156,693 | -0,197 | |  |  |  |  |  |
| ORF6 | SRD06 | estimate | exp | Exponential | 13033,960 | -0,582 | |  |  |  |  |  |
| ORF6 | Yang96 | estimate | log | Exponential | 12954,479 | -0,506 | |  |  |  |  |  |
| ORF6 | Yang96 | estimate | exp | Skyline | 12951,400 | -0,359 | |  |  |  |  |  |

Comparison of alternative Bayesian models used to estimate time-calibrated phylogenies of PRRSV. For each analysis (PRRSV1–PRRSV22 and ORF6), we report the codon or nucleotide substitution model, treatment of base frequencies, molecular-clock parameterisation and coalescent prior, together with the approximate AICM model score (and its standard error) and the mean posterior log-likelihood (with standard deviation, standard error and variance), plus the effective sample size (ESS) of the likelihood. Lower AICM values indicate better model fit. Clock models are denoted as strict, lognormal (log), exponential (exp) or gamma; demographic priors are constant population size (Const), exponential growth (Exponential) or Bayesian skyline (Skyline).
