## Supplementary material for "Rodent Origins and Human-Mediated Evolutionary Dynamics of PRRSV": Table_S4_changes_pig_husbundary.docx

**Table S4 Important changes in pig production management in correlation with PRRS outbreak**

| 1950 | pigs began to be reared more in confined buildings, and the various stages of production became more defined: breeding, gestation, farrowing, lactation, pre-nursery, nursery, grower, and finisher. |
| --- | --- |
| 1960 | in the United Kingdom and about 10 years later in the United States, the pyramid system of producing both gilt and boar breeding stock originated; replacing traditional purebred breeding ever since. which placed a nucleus farm at the apex for development of superior genetic lines. |
| 1975 | The first cryopreserved boar semen has been commercially available in Norway; until the mid-1970s, boars were almost the exclusive means of transfer of genes among farms in the United States. |
| 1979 | Was discovery that the separation of naturally farrowed piglets away from their mothers at weaning would exclude infectious agents led to a profound development in multi-site rearing technique. *Medicated early weaning (MEW)* was coined the term for his discovery because the weaned the pigs early, with heavy medication, into an isolated location away from all other pigs. *Age-segregated rearing (ASR)* fits in this group of MMEW isowean and SEW. They’re all synonyms but are different from MEW. |
| 1988 | Multi- site rearing initiated in the United States in 1988 on a newly constructed 2000-sow farm where the three main stages of pig production were placed on three separate sites was an entirely new concept. Prior to 1988, both finisher pigs and breeding stock were reared either on one-site farrow-to-finish or on traditional two-site production system. In 1989 Harold Trettin in Rockford, Iowa, made the first conversions from a traditional two-site system to a three-site (single source, single locus) farm in 1989. |
| 1988 | The Netherlands, the adoption of AI in pig farming by the late 1960s. Similarly, in other European countries, the adoption of AI in pig farming increased significantly during the 1970s and 1980s, while in 180-90 results of succes of AI imporved tremendously and became advantntageous to use at commercial farms.. https://en.engormix.com/pig-industry/swine-management/artificial-insemination-pigs-research_a40422/?utm |
| 1992 | AI was used several times in the US on commercial sow farms before it became standard practice in the early 1990s. https://www.pigchamp.com/news/benchmark-magazine/articles/ArtMID/2128/ArticleID/197/the-evolution-of-swine-artificial-insemination-2020?utma |
| 1974 | During this period, the Soviet Union intensified efforts to modernize animal husbandry, including pig farming. The focus was on creating large, mechanized farms to increase meat production and meet the growing demand for animal products. https://doi.org/10.12987/yale/9780300200690.003.0004 |
