## Supplementary material for "Rodent Origins and Human-Mediated Evolutionary Dynamics of PRRSV": Table_S5_partions.docx

**Table S5 ORF partitions and genome coordinate ranges by species.**

| Number of sequences | Species | ORF1a | ORF1b | ORF2ab | ORF3 | ORF4 | ORF5 | ORF6 | ORF7 |
| --- | --- | --- | --- | --- | --- | --- | --- | --- | --- |
| 1 | ApRAV1^1^ | 215-6991 | 6979-7048 | 11337-11561 | 12033-12770 | 12668-13210 | 13138-13827 | 13815-14336 | 14333-14737 |
| 1 | DeBMV1^2^ | 210-6464 | 6458-10960 | 10939-11583 | 11454-12095 | 13442-13981 | 13984-14781 | 14739-15260 | 15235-15582 |
| 12 | EAV^3^ | 225-5405 | 5405-9751 | 9751-10507 | 10310-10797 | 10700-11158 | 11146-11913 | 11901-12389 | 12313-12645 |
| 1 | FpgRAV^4^ | 215-6985 | 6985-11358 | 11337-11561 | 12033-12770 | 12668-13210 | 13138-13827 | 13815-14336 | 14333-14737 |
| 1 | FSVV^5^ | 201-6428 | 6422-10819 | 10756-11466 | 12831-13298 | 13133-13645 | 13678-14412 | 14406-14885 | 14866-15201 |
| 1 | KKCBV1^6^ | 154-6321 | 6315-10760 | 10714-11376 | 11250-11858 | 11783-12256 | 13288-14094 | 14088-14573 | 14551-14886 |
| 1 | KRCV1^7^ | 206-6352 | 6346-10740 | 10680-11378 | 11818-12402 | 12309-13047 | 12846-13445 | 13292-13813 | 14582-15067 |
| 1 | KRCV2^8^ | 214-6372 | 6366-10823 | 10706-11377 | 11781-12281 | 12274-12928 | 12808-13386 | 13290-13805 | 13805-14567 |
| 1 | KRTGV1^9^ | 163-6345 | 6339-10814 | 10805-11449 | 11805-12401 | 12394-13606 | 12856-13266 | 13074-13601 | 13604-14389 |
| 1 | LDEV^10^ | 157-6777 | 6850-11046 | 11046-11729 | 11597-12172 | 12064-12591 | 12588-13187 | 13175-13690 | 13677-14024 |
| 1 | MYBV1^11^ | 166-6336 | 6330-10790 | 12279-12509 | 12597-13031 | 12848-13324 | 13317-14087 | 14081-14566 | 14544-14891 |
| 1 | PeV^12^ | 165-6353 | 6347-10822 | 12374-12616 | 12893-13429 | 13270-13800 | 13814-13999 | 14606-15091 | 15066-15413 |
| 37 | PRRSVEU^13^ | 222-7412 | 7394-11785 | 11796-12545 | 12404-13201 | 12946-13497 | 13494-14099 | 14087-14608 | 14598-14984 |
| 38 | PRRSVUS^14^ | 20-7528 | 7528-11901 | 11903-12129 | 12526-13290 | 13071-13607 | 13618-14220 | 14205-14729 | 14719-15090 |
| 2 | RAV^15^ | 3-7415 | 7415-11779 | 11829-12593 | 12607-13182 | 12933-13481 | 13511-14092 | 14077-14601 | 14591-14959 |
| 1 | RoAV^16^ | 1-7572 | 7572-11948 | 11924-12676 | 12535-12144 | 13077-13625 | 13622-14233 | 14218-14742 | 14732-15103 |
| 1 | MgAV1^17^ | 101-7576 | 7702-11934 | 11941-12639 | 12537-12334 | 13211-13762 | 13708-14373 | 14358-14882- | 14872-15240 |
| 1 | FpgRAV^18^ | 215-6985 | 6985-11358 | 11337-11561 | 12033-12770 | 12668-13210 | 13138-13827 | 13815-14336 | 14333-14737 |
| 11 | SHFV^19^ | 210-6521 | 6521-10996 | 10953-11798 | 11486-12100 | 11938-12555 | 14002-14838 | 14832-15320 | 15295-15630 |
| 114 |  |  |  |  |  |  |  |  |  |

Abbreviations: ^1^African pouched rat arterivirus, ^2^DeBrazzas monkey arterivirus, ^3^Equine arterivirus, 4 Forest pouched giant rat arterivirus, ^5^Free State vervet virus, ^6^Kafue kinda chacma baboon virus, ^7^Kibale red colobus virus 1, ^8^Kibale red colobus virus, ^9^Kibale red-tailed guenon virus 1, ^10^Lactate dehydrogenase-elevating virus, ^11^Mikumi yellow baboon virus 1, ^12^Pebjah virus, ^13^PRRSV-1, ^14^PRRSV-2, ^15^Rat arterivirus, ^16^Rodent arterivirus, ^17^Myodes glareolus arterivirus, ^18^Forest pouched giant rat arterivirus, ^19^Simian hemorrhagic fever virus.
